# Early consequences of the phospholamban mutation PLN-R14del^+/-^ in a transgenic mouse model

**DOI:** 10.1101/2023.04.05.535536

**Authors:** Claudia Maniezzi, Marem Eskandr, Chiara Florindi, Mara Ferrandi, Paolo Barassi, Elena Sacco, Valentina Pasquale, Angela S. Maione, Giulio Pompilio, Vivian Oliveira Nunes Teixeira, Rudolf A de Boer, Herman H W Silljé, Francesco Lodola, Antonio Zaza

## Abstract

**Aims:** The heterozygous phospholamban (PLN) mutation R14del (PLN R14del^+/-^) is associated with a severe arrhythmogenic cardiomyopathy (ACM) developing in the adult. “Superinhibition” of SERCA2a by PLN R14del is widely assumed to underlie the pathogenesis, but alternative mechanisms such abnormal energy metabolism have also been reported. This work aims to 1) to evaluate Ca^2+^ dynamics and energy metabolism in a transgenic (TG) mouse model of the mutation prior to cardiomyopathy development; 2) to test whether they are causally connected.

**Methods and Results:** Ca^2+^ dynamics, energy metabolism parameters, reporters of mitochondrial integrity, energy and redox homeostasis were measured in ventricular myocytes of 8-12 weeks-old, phenotypically silent, TG mice. Mutation effects were compared to pharmacological PLN antagonism and analysed during modulation of sarcoplasmic reticulum (SR) and cytosolic Ca^2+^ compartments. Transcripts and proteins of relevant signalling pathways were evaluated. The mutation was characterized by hyperdynamic Ca^2+^ handling, similar to that induced by PLN antagonism. Albeit all components of energy metabolism were depressed at rest, functional signs of mitochondrial damage or energy starvation were absent and cell energy charge was preserved. The response of mitochondrial O_2_ consumption to SERCA2a blockade was lost in mutant myocytes (SR-mitochondrial uncoupling) and ER-stress signalling was activated.

**Conclusions:** 1) PLN R14del^+/-^ loses its ability to inhibit SERCA2a, which argues against SERCA2a superinhibition as a mechanism of ACM; 2) depression of resting energy metabolism may at least partly reflect impairment of SR-mitochondrial coupling; 3) ER-stress may be an early factor in the pathogenesis.

## 1. INTRODUCTION

Phospholamban (PLN) is a small protein which restrains SERCA2a operation, thus limiting Ca^2+^ uptake by the sarcoplasmic reticulum (SR) under resting conditions. Receptor triggered phosphorylation (e.g. by PKA) positively regulates SERCA2a by relieving inhibition by PLN.

The heterozygous deletion of arginine 14 in PLN (PLN R14del^+/-^) is associated with a form of arrhythmogenic dilated cardiomyopathy (ACM), characterized by PLN aggregate formation, myocardial fibrosis, and heart failure, with typical onset at middle age.^1^ PLN R14del^+/-^ is among the prevailing cardiomyopathy-related mutations, particularly in the Netherlands^2^ and currently lacks a specific treatment. More than a decade ago, the pioneering work of Haghighi and coworkers^3^ ascribed PLN R14del^+/-^ phenotype to a “superinhibitory” effect on SERCA2a detected, in a heterologous expression system, as a decrease in sensitivity of pump function to Ca^2+^. Several subsequent studies, inspired by this interpretation, provided more or less indirect support.^4–8^ However, the mechanisms proposed to account for the superinhibitory effect was inconsistent among these studies and remains debated.

The “superinhibition” theory motivated us to test reversal of PLN R14del^+/-^ phenotype by a recently developed compound (PST-3093) that selectively stimulates SERCA2a by antagonizing its interaction with PLN.^9^ Strikingly, human iPS-derived cardiomyocytes (hiPS-CMs) from a heterozygous human mutation carrier displayed a hyperdynamic Ca^2+^ handling instead, a phenotype that obviously is incompatible with the SERCA2a “superinhibition”. Notably, PST-3093 mimicked the effect of the mutation when applied to the WT hiPS-CMs, but it was ineffective in mutant ones.^10^ Studies on contracting engineered tissues (EHT), obtained from the same hiPS-CMs, detected a major decrease in force development and energy metabolism derangements, but no clear abnormalities in intracellular Ca^2+^ dynamics.^11^ Nonetheless, being such studies based on immature cells from a single mutation carrier, caution clearly is warranted in generalization of these results, particularly in light of the wide acceptance of the superinhibition hypothesis.

We have developed and deeply characterized a PLN R14del^+/-^ transgenic (TG) mouse model which closely recapitulates the human ACM phenotype, including lack of cardiac abnormalities at young age.^12^ The present work characterizes cardiomyocytes (CMs) obtained from this disease model in terms of mutation effect on intracellular Ca^2+^ dynamics and energy metabolism. This with the aim of testing the “superinhibition” theory in native mature CMs and compare them to hiPS-CMs for further functional abnormalities contributing to ACM pathogenesis, thus providing cross-validation of experimental models.

Evaluation of primary pathogenetic mechanisms in mutations leading to contractile deficit is complicated by overlap with the etiology-unspecific maladaptive process induced by the deficit itself, in which SERCA2a loss of function is conspicuous.^13^ To minimize this potential confounder, we selected an animal age at which cardiac contraction is still normal in the PLN R14del^+/-^ TG mice.^12^

## 2. MATERIALS AND METHODS

The methods are described here to the extent of allowing interpretation of results. A detailed description of the methods is given in the Supplement.

### 2.1 Experimental model

The studies were performed on tissues from PLN R14del^+/-^ (Mut) mice aged 8 to 12 weeks and their WT littermates (controls). In this age range, the animals are healthy; overt physical or echocardiographic signs of chamber remodelling or contractile dysfunction are present in this model at 18 months of age.^12^ This is crucial to the interpretation of the observed changes as directly resulting from the mutation, rather than secondary to aspecific myocardial remodelling.

Myocytes were enzymatically dissociated with a manual perfusion method,^14^ which does not discriminate between right and left ventricles and studied within 24 hours.

All experiments involving animals confirmed to the guidelines for Animal Care endorsed by the Milano-Bicocca and to the Directive 2010/63/EU of the European Parliament on the protection of animals used for scientific purposes.

### 2.2 Measurements and techniques

- SERCA2a ATPase activity was measured in myocardial homogenates at multiple Ca^2+^ concentrations; data points were fitted to a sigmoidal (Hill) function, from which the maximum velocity (V_max_) and Ca^2+^ affinity [Kd_Ca_] parameters were estimated.
- Electrophysiological experiments were carried out by whole-cell patch clamp on isolated cardiomyocytes (CMs) in the ruptured- or perforated-patch configuration, as specified in the relevant results sections.
- Cytosolic Ca^2+^ was optically measured from isolated CMs by using Fluo 4-AM as the Ca^2+^ – sensitive probe; fluorescence (F) was normalized to the value measured after prolonged quiescence (F0). Ca^2+^ measurements were carried out in field-stimulated CMs, or V-clamped CMs, as specified in the relevant results sections.
- Oxygen consumption rate (OCR, an index of oxidative phosphorylation) and Proton Efflux Rate (PER, an index of anaerobic glycolysis) were measured on quiescent CMs seeded in multiwell plates using the Seahorse technology (Agilent Extracellular Flux Analyzer XFe96). All measurements were normalized to the (automatically counted) number of viable CMs in the measuring well. All the experimental conditions were represented in each plate to allow comparisons within the same experiment; the position of each condition in the plate was swapped between experiments to prevent technical bias. The preparations were exposed to reagents to obtain specific OCR and PER parameters as described in the Experimental Protocols section.
- Radical Oxygen Species (ROS) content of CMs was measured after incubation with the fluorescent probe 2’7’-dichclorofluorescin diacetate (DCFDA). Automated (unbiased) confocal single-cell fluorescence measurement was performed by Operetta CLS™ (Operetta– Perkin Elmer) at 40× magnification.
- Mitochondrial membrane potential (Ψ_m_) was measured from CMs using Tetramethylrhodamine, Ethyl Ester (TMRE) as probe. Stained CMs were seeded at the concentration 2.5×10^3^ per well. TMRE fluorescence was analysed by confocal imaging at 63× magnification (Operetta CLS™). At the concentration used in the present study, TMRE works in “quenching mode”, i.e. emission increases as Ψ_m_ depolarizes.^15^ At any rate, TMRE fluorescence signal was calibrated in each measurement by short-circuiting mitochondrial electron transport with FCCP.
- qRT-PCR was performed on mRNA extracted from myocardial samples and quantified by a NanoDrop spectrophotometer. qRT-PCR was performed with the primers reported in Supplement Table 1. All reactions were performed in a 384-well format. The relative quantities of specific mRNAs were obtained by the delta-delta Ct method with normalization to the housekeeping gene glyceraldehyde 3-phosphate dehydrogenase (GAPDH).
- Western Blot Analysis was performed on RV samples. Total protein extracts were subjected to SDS-PAGE and transferred onto a nitrocellulose membrane, blocked for 1 hour at room temperature and incubated overnight at 4°C with the appropriate primary antibodies (reported in Supplement Table 2). Peroxidase-conjugated secondary antibodies were then applied for 1 hour and the peroxidase signal visualized using a chemiluminescent substrate. Blot images were acquired and blot densitometric analysis was performed by ImageJ software. Protein signals were normalized to GAPDH or TUBULIN ones.

### 2.3 Experimental protocols

Significant parameters were extracted from the above measurements by applying suitable experimental protocols.

- Ca^2+^ transients (CaT) were evaluated in intact CMs., field-stimulated at 1 Hz. Ca^2+^ transient amplitude (CaT amplitude), Ca^2+^ transient decay kinetics (τ _decay_), Ca^2+^ transient rise-time (t_peak_) and diastolic Ca^2+^ (CaD) were measured. Rate-dependency of CaT properties was tested by stepwise increments in pacing rate (to 1, 1.3, 1.7, and 2 Hz).
- Sarcoplasmic reticulum (SR) Ca^2+^ content was estimated in V-clamped CMs as the integral of the I_m_ (mostly representing I_NCX_) elicited by a caffeine (10 mM) pulse, applied after a loading train of V steps (−40 to 0 mV at 1 Hz).^16^ To avoid extracellular Ca^2+^ influx, caffeine was dissolved in Ca^2+^-free solution (containing 1 mM EGTA CsOH).
- The “gain” of Ca^2+^-induced Ca^2+^ release (CICR) was measured in V-clamped CMs as the Ca^2+^ release/influx ratio, as previously described.^17^
- The fraction of SR Ca^2+^ content released by membrane excitation (fractional release) was calculated as the ratio between the amplitudes of V- and caffeine-triggered CaT.
- Information on NCX function was obtained by linear fitting of the trailing branch of the I_NCX_/[Ca^2+^] loops, recorded during caffeine-induced transients. The slope coefficient and the 0 I_NCX_ intercept were used as surrogate of NCX “conductance” and cytosolic [Ca^2+^] at electrochemical equilibrium, respectively.
- The Ca^2+^ uptake function of the SR was evaluated through a “SR reloading” protocol, applied under V-clamp; CaT and I_CaL_ were simultaneously recorded.^18^ The SR was initially emptied by a caffeine pulse and progressively reloaded by a train (0.25 Hz) of 200 ms V steps from −40 to 0 mV. The following parameters were analysed from each step of the protocol: i) CaT amplitude, ii) CICR gain, iii) τ _decay_, iv) CaD. The rate of increment of the former two parameters during the loading protocol reports the rate of SR refilling. The time constant of CaT decay (τ _decay_) reports the rate of cytosolic Ca^2+^ clearance (the faster Ca^2+^ removal, the smaller τ _decay_) within each step, i.e. at varying SR filling levels. CaD course reports the rate of cytosolic Ca^2+^ accumulation.
- OCR and PER parameters were obtained by sequential exposure of preparations to Oligomycin A (ATP synthase inhibitor), FCCP (in OCR only, short-circuits the electron transfer chain, ETC), Rotenone/Antimycin (block ETC complexes I and III) or 2-deoxy-D-glucose (2DG in PER only, inhibits glycolysis) as follows:
  `o` OCR: i) mitochondrial basal respiration = OCR before Oligomycin – OCR after Rotenone/Antimycin; ii) mitochondrial maximal respiration = OCR after FCCP − OCR after Rotenone/Antimycin; iii) spare respiratory capacity = mitochondrial maximal respiration – mitochondrial basal respiration; iv) non-mitochondrial respiration = OCR after Rotenone/Antimycin.
  `o` PER: i) basal glycolysis = PER before Oligomycin – PER after 2DG; ii) compensatory glycolysis = PER after Rotenone/Antimycin − PER after 2DG; iii) glycolytic reserve = compensatory glycolysis – basal glycolysis; iv) non-glycolytic acidification = PER after 2DG.
- I_KATP_ was measured as a surrogate reporter of the ADP/ATP ratio. CMs were clamped at – 120 mV in the perforated-patch configuration; I_KATP_ was measured as glibenclamide-sensitive current.

### 2.4 Statistical analysis

Statistical analysis was carried out with GraphPad Prism 8. Normality of distribution was assessed using D’Agostino-Pearson’s normality test. Comparison of sample means was carried out with parametric or non-parametric tests, according to the data type (continuous or categorical). Parametric or non-parametric ANOVA (with the respective post-hoc corrections) were used for multiple comparisons of continuous or categorical data respectively. In the case of repeated measurements (e.g. rate-dependency and SR loading protocols) a mixed-effects ANOVA model containing “Treatment” (WT vs Mut, PST-3093 vs Control) and “Variable” (rate or step #) factors was used. Significance of “Treatment X Variable interaction” (i.e. difference between Treatments in their response to the Variable) was first tested; in its absence, significance of difference between Treatments at all Variable values was tested. In figures, whenever feasible, individual data points were plotted, along with the sample mean ± SEM, to illustrate dispersion. Whenever the threshold for statistical significance (p<0.05) was achieved, the actual p-value for the comparison was reported as an index of robustness. For each experiment, the number of preparations or cells (n) and the number of animals from which they were obtained (N) are indicated in the respective figure legend.

## 3. RESULTS

### 3.1 SERCA2a ATPase activity

SERCA2a ATPase activity correlates with Ca^2+^ transport rate;^19^ its measurement in myocardial homogenates provides direct information on the transporter function in a simplified (cell-free) system. Ca^2+^-sensitivity of SERCA2a activity is mostly determined by SERCA2a-PLN interaction and is therefore suitable to detect its abnormalities.^19^

In myocardial homogenate preparations, the Ca^2+^ dissociation constant [Kd_Ca_] of SERCA2a ATPase activity was 21% lower (unpaired Student’s t-test, p=0.0005) in Mut; the maximum velocity of SERCA2a ATPase activity was similar between Mut and WT preparations. Mut SERCA2a ATPase activity exceeded WT one above 300 nM Ca^2+^ (**Figure 1**).

**Figure 1.**
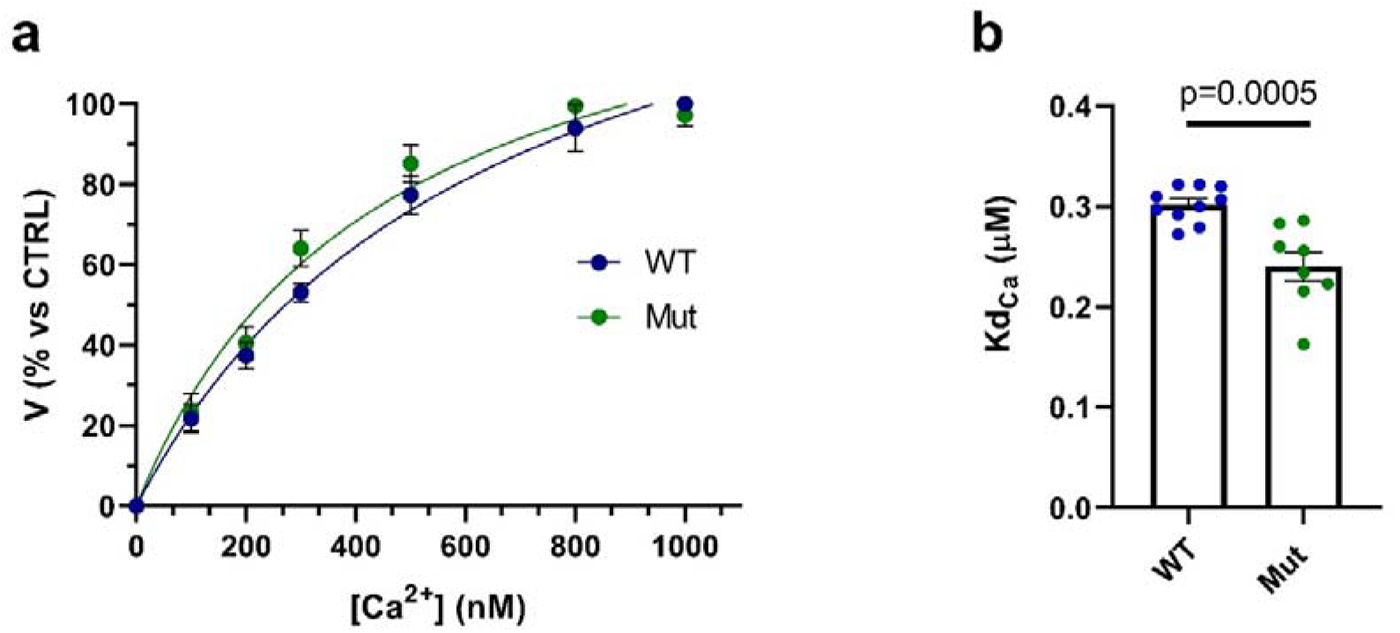
Ca^2+^-dependent activation of SERCA2a ATPase in WT and Mut cardiac homogenates. (**a**) Experimental points fitted by Hill functions. (**b**) Affinity constant for Ca^2+^-dependent activation (Kd_Ca_) estimated from the fitting (WT: 0.30 ± 0.01, n=10; Mut: 0.24 ± 0.01, n=8). WT: N=2; Mut: N=2. Data are expressed as mean ± SEM. Unpaired Student’s t-test.

This observation suggests reduced inhibition of SERCA2a by PLN in Mut preparations.

### 3.2 Intracellular Ca^2+^ dynamics

Next we evaluated SERCA2a function in the context of an intact myocyte. To improve mechanistic interpretation of the mutation effect, studies with an agent known to increase SERCA2a function by preventing its interaction with PLN^9^ were included. SERCA2a function may have different impact on intracellular Ca^2+^ dynamics at different heart rates; therefore, the rate-dependency of the variable’s (Mut vs WT or PST-3093 vs Control) effect on Ca^2+^ dynamics was also evaluated in a separate set of CMs, field-stimulated at four rates between 1 and 2 Hz. Since these experiments aim to compare the rate-dependency of effect, statistical significance refers to the Treatment X Rate interaction.

#### 3.2.1 Effect of the mutation

##### Steady-state stimulation (1 Hz)

Mut effect on intracellular Ca^2+^ dynamics was assessed in intact WT and Mut CMs during steady-state field-stimulation at 1 Hz (**Figure 2**). Ca^2+^ transient (CaT) parameters were measured as described in methods.

**Figure 2.**
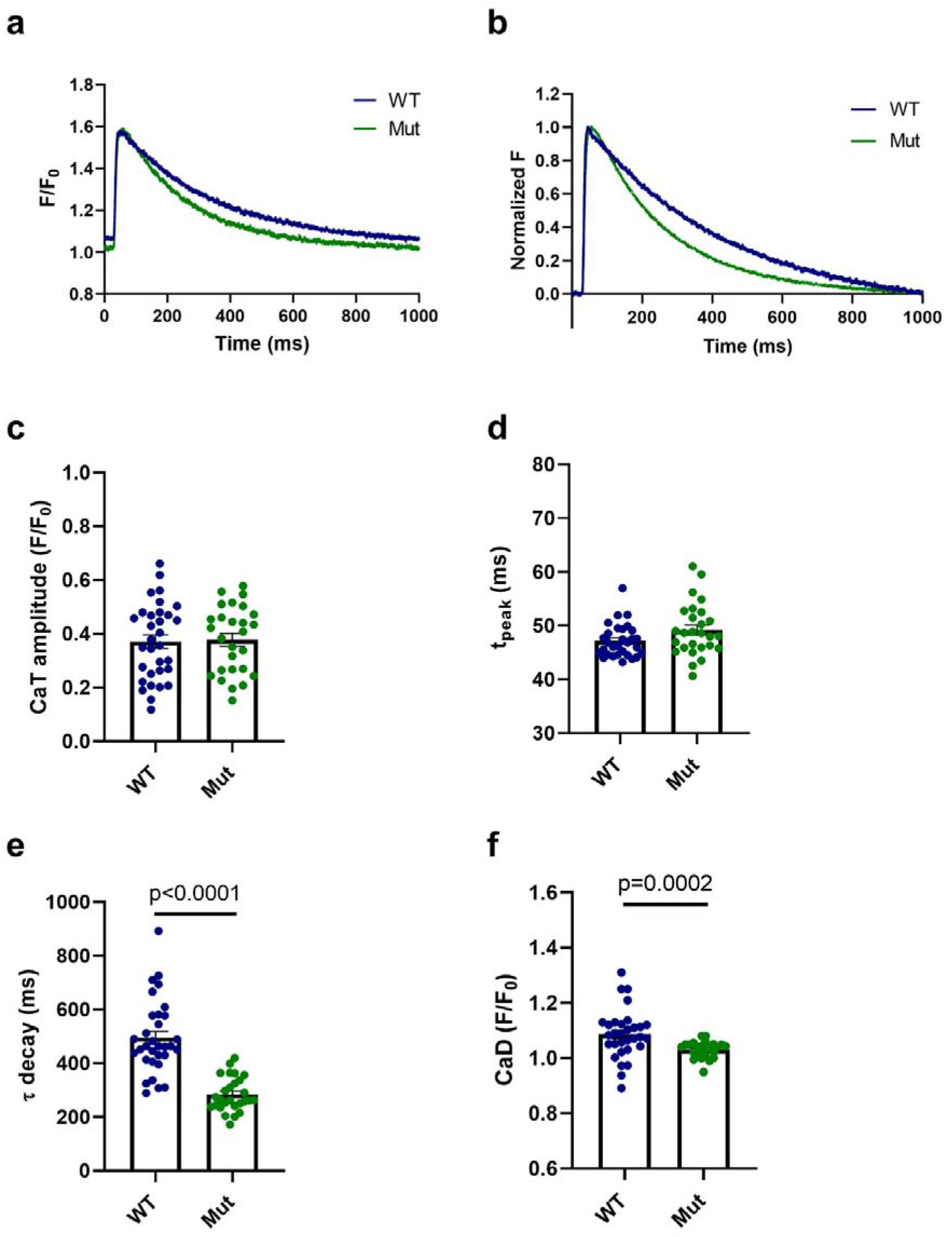
Parameters of calcium transients in WT and MUT. **(a)** Representative Ca^2+^ transients (CaT). (**b**) Normalized CaT traces. (**c**) CaT amplitude (WT: 0.37 ± 0.03, n=32; Mut: 0.38 ± 0.02, n=27). (**d**) CaT time to peak, t_peak_ (WT: 47.1 ± 0.56, n=30; Mut: 49.2 ± 0.94 n=27). (**e**) Ca^2+^ transient decay kinetics, τ _decay_ (WT: 495 ± 24.1, n=32; Mut 284 ± 12.2, n=27). (**f**) Diastolic Ca^2+^, CaD (WT: 1.09 ± 0.02, n=31; Mut: 1.03 ± 0.01, n=27). WT: N=4; Mut: N=4. Data are expressed as mean ± SEM. Mann-Whitney U-test.

In Mut CMs τ _decay_ (**Figure 2e**) was reduced by 43% (Mann-Whitney U-test, p<0.0001 vs WT); diastolic Ca^2+^ (CaD, **Figure 2f**) was 5% lower (Mann-Whitney U-test, p=0.0002 vs WT) and remarkably less variable. CaT amplitude (**Figure 2c**) and rise-time (t_peak,_ **Figure 2d**) were comparable between the two genotypes.

##### Rate-dependency

The following CaT parameters showed significant rate-dependency in WT CMs (**Figure S1**): t_peak_ (inverse, p<0.0001), τ _decay_ (inverse, p<0.0001) CaD (direct, p<0.0001).

As compared to WT, in Mut CMs: τ _decay_ had a shallower rate-dependency (**Figure S1c**, Mixed-effects model, Treatment X Rate, p=0.0015 vs WT) due to preferential shortening at slow rates. As expected from faster SR Ca^2+^ uptake, CaD accumulation was less pronounced in Mut CMs (**Figure S1d**, Mixed-effects model, Treatment X Rate, p=0.0114 vs WT).

For the remaining parameters rate-dependency was similar between the two genotypes.

##### SR reloading rate and steady-state SR Ca^2+^ content

SR Ca^2+^ uptake function was assessed in V-clamped CMs by the “SR reloading” protocol and by measuring caffeine-releasable SR Ca^2+^ content (CaSR) (**Figure 3a**, see methods). The difference between Mut and WT CMs in the behaviour of CaT parameters during SR reloading (from step 1 through 15) is reported below (**Figure 3**).

**Figure 3.**
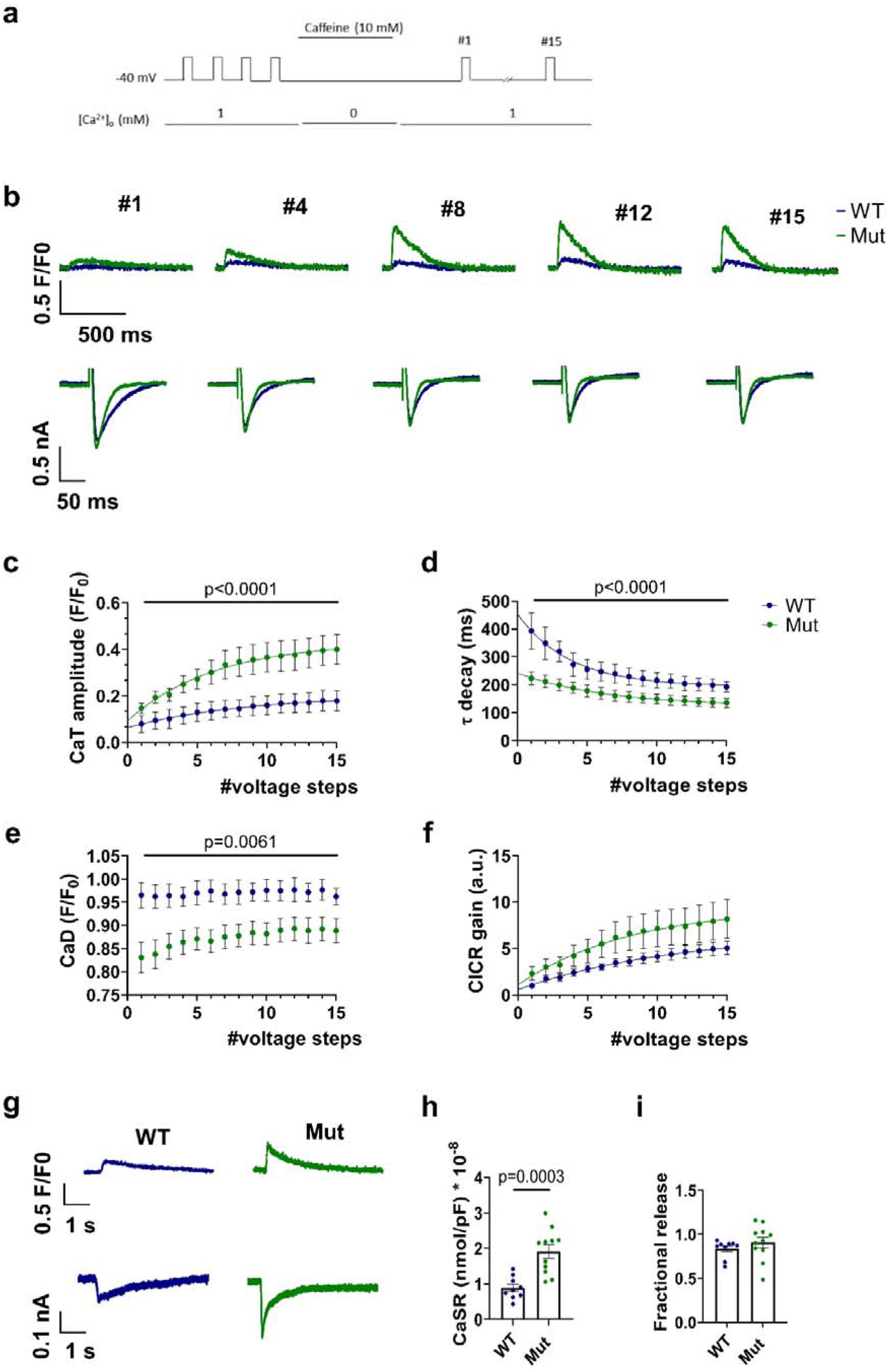
SR Ca^2+^ content and uptake in WT and Mut. (**a**) Experimental protocol. (**b**) Representative traces of Ca^2+^ transients (upper panels) and voltage-dependent calcium currents (I_CaL_) (lower panels) recorded during SR reloading after caffeine-induced depletion. CaT baselines are superimposed to emphasize changes in transient amplitude. (**c-f**) Mean values of CaT parameters measured during each voltage step (1–15) of the loading train. WT: n=9, N=7; Mut: n=11, N=6. Mixed-effects model for loading. (**g**) Representative traces of caffeine-induced Ca^2+^ Transients (upper panels) and membrane currents, I_m_ (lower panels) recorded during caffeine-induced SR depletion. (**h**) SR Ca^2+^ content (CaSR) measured from the integral of I_m_, reflecting Ca^2+^ efflux through the Na^+^/Ca^2+^ exchanger, I_NCX_ (WT: 0.89 ± 0.11, n=9; Mut: 1.91 ± 0.19, n=11). (**i**) Fractional release (WT: 0.83 ± 0.04, n=9; Mut: 0.91 ± 0.06, n=11). WT: N=7; Mut: N=6. Data are expressed as mean ± SEM. Unpaired Student’s t-test.

CaT amplitude progressively increased during SR reloading, as expected (**Figure 3c**). The rate of such increment, as well as the final amplitude value, were larger for Mut CMs (Mixed-effects model, Treatment X Step, p<0.0001 vs WT). The same pattern was observed after CaT normalization for Ca^2+^ influx through I_CaL_ (**Figure 3f**, CICR gain); however, due to the larger variance in Mut data, for this parameter statistical significance was not achieved.

The τ _decay_ progressively decreased during SR reloading (**Figure 3d**). In Mut CMs it started off from considerably lower values, but its progressive decrement over the protocol was shallower (Mixed-effects model, Treatment X Step, p<0.0001 vs WT). CaD started off from a lower value in Mut CMs (unpaired Student’s t-test, p=0.0060), but the difference decreased thereafter (**Figure 3e**, Mixed-effects model, Treatment X Step, p=0.0061 vs WT).

Under the conditions of this experiment, SR Ca^2+^ content (CaSR) was considerably larger in Mut CMs (**Figure 3h**, unpaired Student’s t-test, p=0.0003 vs WT); fractional release was similar between Mut and WT CMs (**Figure 3i**).

*To summarize, as compared to WT ones, Mut CMs were characterized by “hyperdynamic” Ca^2+^ handling, compatible with SERCA2a enhancement (as opposed to inhibition). This pattern was consistent between CaT parameters of intact CMs during field-stimulation and evaluation of SR reloading under V-clamp.*

##### NCX “conductance” and equilibrium

NCX functional parameters were measured during the caffeine pulse, i.e., under conditions short-circuiting SR transports (**Figure S2**).

In Mut CMs NCX “conductance” was significantly decreased (**Figure S2b**, Mann-Whitney U-test, p=0.02 vs WT); Ca^2+^ concentration at the (extrapolated) 0 I_NCX_ value (transport equilibrium point) was not significantly changed (**Figure S2c**).

#### 3.2.2 Effect of PLN antagonism (PST-3093) in MUT vs WT CMs

##### Steady state stimulation (1 Hz)

The effect of PLN antagonism by PST-3093 on intracellular Ca^2+^ dynamics was assessed in intact WT and Mut CMs during steady-state field-stimulation at 1 Hz. CaT parameters were measured as described in the methods section.

In WT CMs, PST-3093 reduced τ _decay_ by 24% (**Figure S3e**, One-way ANOVA, post-hoc p=0.0124 vs Control); the remaining parameters were unaffected (**Figure S3**).

In Mut CMs, PST-3093 failed to affect the CaT parameters significantly (**Figure S3**), notably including τ _decay_ (**Figure S3e**).

##### Rate-dependency

For both WT and Mut CMs, CaT parameters showing significant rate-dependency (direct or inverse) under control conditions were: t_peak_ (inverse, p<0.0001), τ _decay_ (inverse, p<0.0001) and CaD (direct, p<0.0001) (**Figures S4 and S5**).

In Mut CMs, PST-3093 enhanced rate-dependent accumulation of CaD (**Figure S4d**, Mixed-effects model, Treatment X Rate, p=0.004 vs Control).

Rate-dependency of all the remaining parameters was not significantly affected by PST-3093 in both genotypes (**Figures S4 and S5**).

*To summarize, in WT CMs PST-3093 had, as postulated, effects compatible with SERCA2a activation (hyperdynamic Ca^2+^ handling), although quantitatively smaller than those exerted by the mutation. On the other hand, PST-3093 effects were nil, or even opposite, in Mut CMs.*

### 3.3 Energy metabolism

Derangement of mitochondrial function is a common feature of the (secondary) “remodelling” process, is related to mishandling of intracellular Ca^2+^ and it ultimately contributes to the development of heart failure. The aim of this set of experiments was to test whether the energy metabolism was altered in Mut CMs at a stage preceding overt contractile failure, thus possibly acting as a primary pathogenetic factor instead. Notably, because of technical constraints, metabolic measurements were performed in unstimulated (quiescent) CMs.

#### 3.3.1 Oxygen consumption rate

O_2_ consumption rate (OCR) was compared between quiescent WT and Mut CMs within the same multiwell plate (**Figure 4**). OCR values were normalized for the number of cells in each well. Mitochondrial respiration profile, from which the respiration parameters were derived, was obtained by modulating specific functions with pharmacological agents, as described in the methods section. The pattern observed in Mut CMs is reported below as relative to that of WT ones.

**Figure 4.**
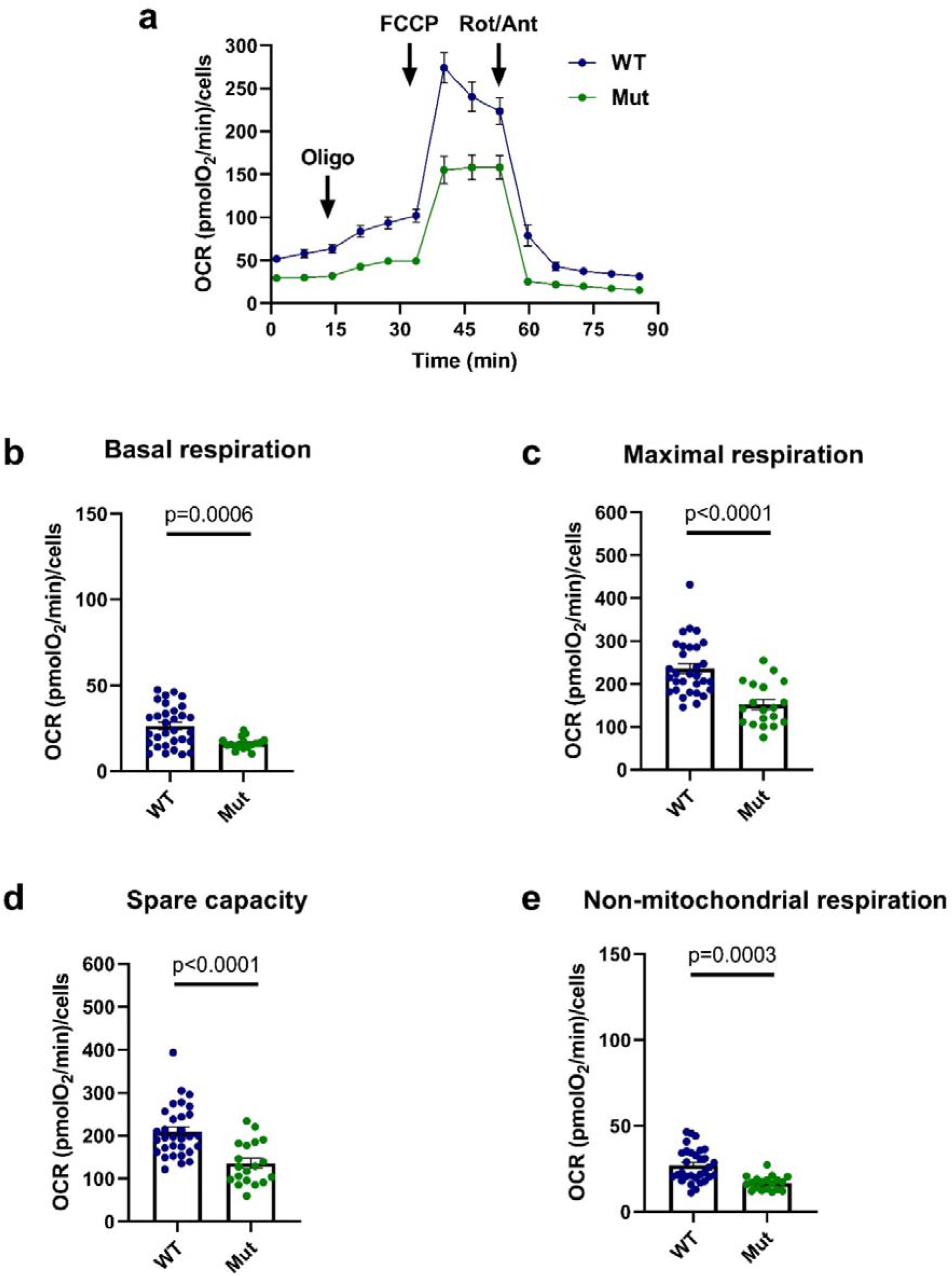
Parameters of mitochondrial respiration in WT and Mut. (**a**) Oxygen Consumption Rate (OCR) profiles of WT and Mut CMs subjected to sequential injections of 1.5 µM Oligo, 0.5 µM FCCP and 3 µM Rot/Ant A (XF Mito Stress Test protocol). (**b**) Mitochondrial basal respiration (WT: 26.40 ± 2.13, n=31; Mut: 16.20 ± 0.76, n=19). (**c**) Mitochondrial maximal respiration (WT: 235 ± 11.46, n=31; Mut: 151 ± 11.54, n=19). (**d**) Spare respiratory capacity (209 ± 10.76, n=31; Mut: 129 ± 12.6, n=17). (**e**) Non-mitochondrial respiration (WT: 26.97 ± 1.74, n=31; Mut: 16.68 ± 0.94 n=19). WT: N=3; Mut: N=3. Data are expressed as mean ± SEM. Unpaired Student’s t-test and Mann-Whitney U-test.

Overall, OCR was significantly depressed in Mut CMs (**Figure 4a**); the observation was reproduced with remarkable consistency across all the preparations tested. The largest difference was found in the mitochondrial maximal respiration (**Figure 4c**, -36%; Mann-Whitney U-test, p<0.0001); nonetheless, mitochondrial basal respiration (**Figure 4b**, -39%; unpaired Student’s t-test, p=0.0006) and spare respiratory capacity (**Figure 4d**, -35%; Mann-Whitney U-test, p<0.0001) were also reduced. Notably, significantly lower values were also found for the non-mitochondrial component of OCR (**Figure 4e**, -38%; unpaired Student’s t-test, p=0.0003).

Modulation of OCR by cytosolic Ca^2+^, and by the SR compartment specifically, was evaluated in a subset of samples. To this end, cytosolic Ca^2+^ was chelated by BAPTA-AM (BAPTA) and SERCA2a was inhibited by Thapsigargin (THAPSI) (**Figure S6**). BAPTA affected OCR parameters in neither WT nor Mut preparations. THAPSI significantly decreased mitochondrial basal respiration (**Figure S6a**; -54% vs control, Kruskal-Wallis, p=0.0265) in WT preparations only and without affecting mitochondrial maximal respiration.

#### 3.3.2 Glycolysis

Anaerobic glycolytic metabolism was evaluated, in the same plates subjected to OCR measurements, as proton efflux rate (PER) (**Figure 5**). PER values were normalized for the number of cells in the wells of the Seahorse plate. PER was measured under basal conditions, after inhibiting mitochondrial respiration (by Rotenone/Antimycin A) to assess compensatory glycolysis, and after blocking glycolysis by 2-deoxy-D-glucose (2DG) to assess non-glycolytic acidification.

**Figure 5.**
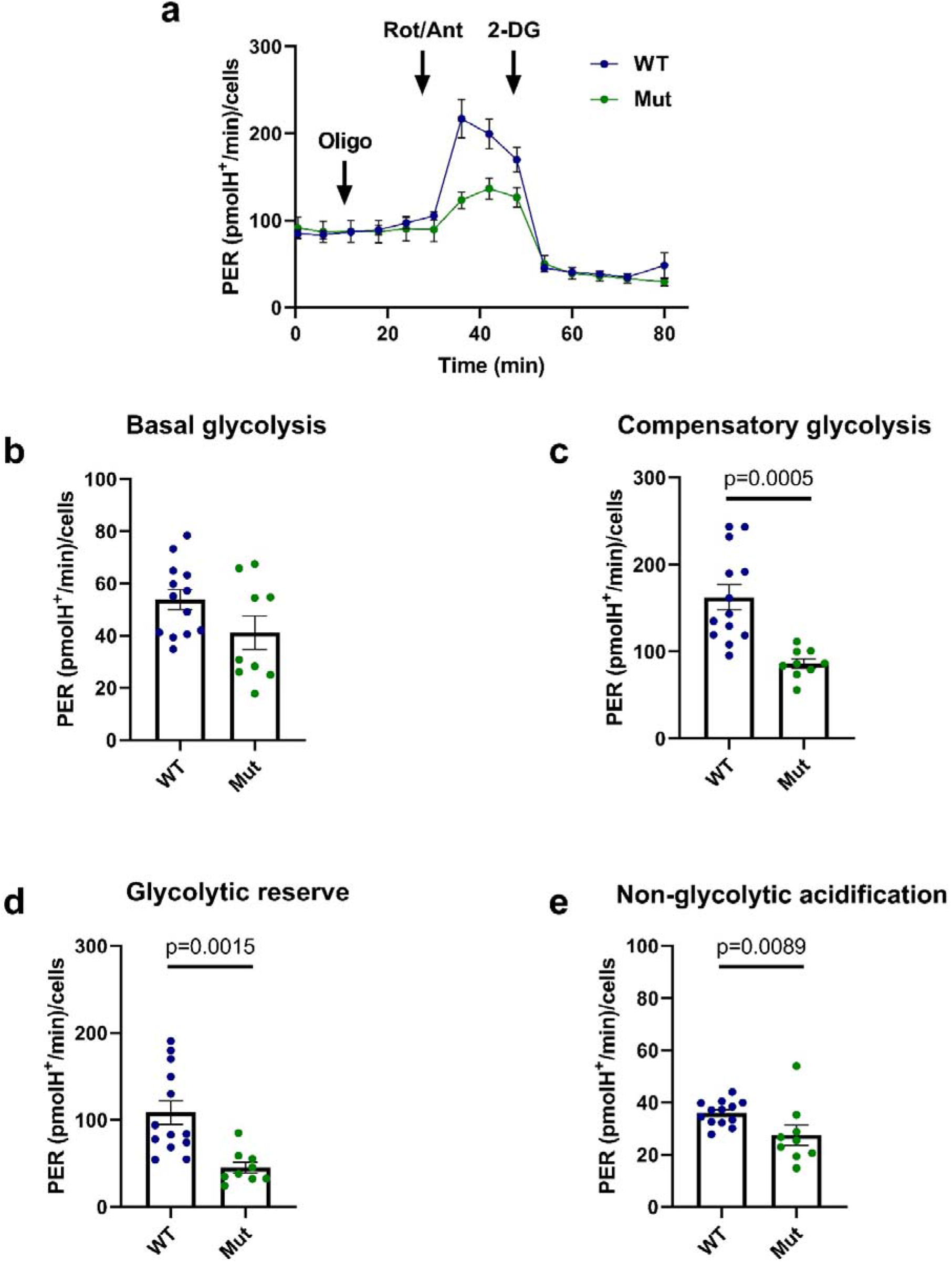
Parameters of anaerobic glycolysis in WT and Mut. (**a**) Proton Efflux Rate (PER) profile of WT and Mut CMs subjected to sequential injections of 1,5 µM Oligo, 3 µM Rot/Ant and 50 mM 2-DG. (**b**) Basal glycolysis (WT: 53.77 ± 3.84, n=13; Mut: 41.15 ± 6.42, n=9). (**c**) Compensatory glycolysis (WT: 162 ± 14.52, n=13; Mut: 86.34 ± 5.48, n=9). (**d**) Glycolytic reserve (WT: 109 ± 13.58, n=13; Mut: 45.19 ± 6.23, n=9). (**e**) Non-glycolytic acidification (WT: 36 ± 1.29, n=13; Mut: 27.56 ± 3.84, n=9). WT: N=3; Mut: N=3. Data are expressed as mean ± SEM. Unpaired Student’s t-test and Mann-Whitney U-test.

Anaerobic glycolysis was depressed in Mut CMs as compared to WT ones (**Figure 5a**). While basal glycolysis showed only a trend to reduction (**Figure 5b**, -23%; unpaired t-test, p=0.0882), compensatory glycolysis (**Figure 5c**, -47%; unpaired Student’s t-test, p=0.0005) and glycolytic reserve (**Figure 5d**, -58%; unpaired Student’s t-test, p=0.0015) were severely depressed. Non-glycolytic acidification was more slightly (-23%), but still significantly, reduced (**Figure 5e**, Mann-Whitney U-test, p=0.0089).

#### 3.3.3 Intracellular ROS and mitochondrial membrane potential (**Ψ**_m_)

Intracellular ROS content was estimated in quiescent CMs by measuring DCFDA fluorescence; confocal images were automatically analysed to quantify the signal from individual vital cells. In Mut CMs, DCFDA signal was marginally, but significantly, lower than in WT CMs (**Figure S7a**, – 5%; unpaired Student’s t-test, p=0.0263 vs WT), to indicate a slight reduction in ROS content.

Mitochondrial membrane potential (Ψ_m_) was evaluated by the fluorescent probe TMRE. TMRE signal was similar between Mut and WT CMs (**Figure S7b**). In the same CMs of both groups, short-circuit of mitochondrial membrane by FCCP significantly increased TMRE signal, thus confirming responsiveness of the probe to Ψ_m_ changes (**Figure S7b-inset).**

#### 3.3.4 Cell energy charge

To assess whether the reduction in energy metabolism of Mut CMs leads to energy starvation, we measured the current carried by the glibenclamide-sensitive K^+^ channel (I_KATP_), whose conductance is steeply proportional to the ADP/ATP ratio.^20^ This surrogate measurement was adopted because, in preliminary experiments, we found bulk fluorescence ATP assays to be confounded by inconstant viability of CMs in the preparations. To avoid intracellular dialysis by the pipette content, measurements were performed with the perforated-patch technique.

I_KATP_ was recorded as glibenclamide-sensitive current at a holding potential of -120 mV, at which K^+^ currents are expectedly inward (i.e., a positive current shift reflects a decrease in I_KATP_). I_KATP_ was normalized to membrane capacitance to obtain current density.

No significant difference in the mean I_KATP_ density was observed between WT and Mut CMs nonetheless, very large I_KATP_ values were occasionally recorded in Mut CMs (**Figure S8**).

*To summarize, both the oxidative and anaerobic components of energy metabolism were depressed in Mut CMs. Nonetheless, the cell energy charge was not decreased, and functional signs of mitochondrial damage were absent. The effect of SERCA2a blockade on OCR was lost in the mutant, thus suggesting impaired Ca^2+^ transfer between the SR and mitochondrial compartments.*

### 3.4 Transcript analysis

The functional derangements reported above may suggest profound changes in Mut CMs biology, may they depend on altered Ca^2+^ handling, or on direct toxicity of mutated PLN. In a broad, but focused, attempt to identify specific processes involved in the overall cell response to the mutation, we analysed transcriptional expression of elements clustering in the following functional groups: 1) remodelling of mitochondria (Mitophagy) and their SR contact sites (MERCS); 2) antioxidant response (DETOX); 3) response to energy starvation (AMPK); 4) response to ER stress (UPR); 5)

CaMKII signalling (**Figure S9**). Western blots were obtained for pErk/Erk (**Figure 6a**), Bax (**Figure 6b**), IP_3_R, pAMPK/AMPK (**Figure 6c**), and pCaMKII/CaMKII (**Figure 6d**).

**Figure 6.**
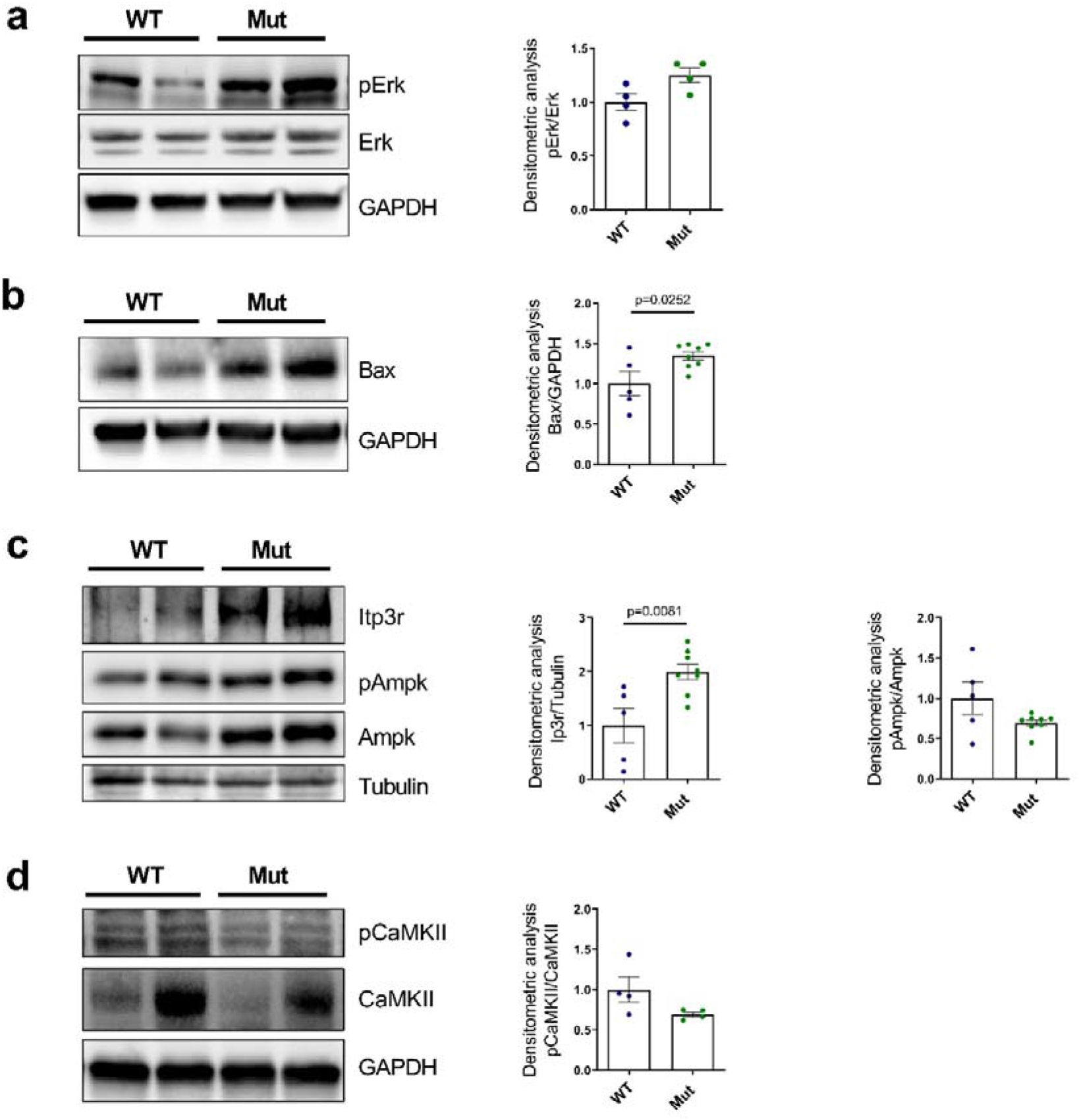
Protein analysis in WT and Mut. (**a**) left: representative images of WB analysis of proteins extracted from RV bioptic samples from WT and Mut mice, hybridized with anti-pErk and anti-Erk antibodies. Immunostaining of the housekeeping GAPDH is included to show total protein load. Right: data from immunoblots were quantified by densitometric analysis: pErk levels were corrected by total Erk densitometry (WT: 1.0000 ± 0.0777, N=4; Mut: 1.2493 ± 0.0691, N=4). (**b**) left: representative images of WB analysis of proteins extracted from RV bioptic samples from WT and Mut mice, hybridized with anti-Bax antibody. Immunostaining of the housekeeping GAPDH is included to show total protein load. Right: data from immunoblots were quantified by densitometric analysis: Bax levels were corrected by GAPDH densitometry (WT: 1.0000 ± 0.1506, N=5; Mut: 1.3475 ± 0.0529, N=8). (**c**) left: representative images of WB analysis of proteins extracted from RV bioptic samples from WT and Mut mice, hybridized with anti-Ip3r, anti-pAmpk and anti-Ampk antibodies. Immunostaining of the housekeeping Tubulin is included to show total protein load. Right: data from immunoblots were quantified by densitometric analysis: Ip3r levels were corrected by Tubulin densitometry (WT: 1.0000 ± 0.3180, N=5; Mut: 1.9938 ± 0.1457; N=8) while pAmpk levels were corrected by total Ampk densitometry (WT: 1.0000 ± 0.2013, N=5; Mut: 0.6950 ± 0.0396, N=8). (**d**) left: representative images of WB analysis of proteins extracted from RV bioptic samples from WT and Mut mice, hybridized with anti-pCaMKII and anti-CaMKII antibodies. Immunostaining of the housekeeping GAPDH is included to show total protein load. Right: data from immunoblots were quantified by densitometric analysis: pCaMKII levels were corrected by total CaMKII densitometry (WT: 1.0000 ± 0.1566, N=4; Mut: 0.6936 ± 0.0309, N=4). Data are expressed as mean ± SEM. Unpaired Student’s t-test.

The transcription of genes involved in UPR and mitophagy were unchanged (**Figure S9**). Nonetheless, a trend to activation of signals involved in the response to ER-stress (pErk/Erk and Bax) was observed at the protein level (**Figure 6a and 6b**).

Genes encoding a couple of AMPK isoforms were upregulated by 1.5-2 fold (**Figure S9**); however, the pAMPK/AMPK ratio showed, if anything, a trend to decrease, thus arguing against activation of energy starvation signalling (**Figure 6c**).

Genes encoding IP_3_R were unchanged (**Figure S9)**; nonetheless IP_3_R protein level was significantly increased (**Figure 6c**), perhaps as a compensation to derangement of SR-mitochondrial communication.^11^

Genes encoding CaMKII were unchanged; moreover, at protein level, the pCaMKII/CaMKII ratio was not increased (**Figure 6d**). Thus, CaMKII signalling may not be activated at this disease stage.

*To summarize, transcript and protein analysis mainly led to negative conclusions regarding activation of energy starvation, CaMKII signalling or ROS detoxification. The few positive results might hint to activation of ER-stress response and impaired SR-mitochondria interaction*.

## 4. DISCUSSION

The main findings of this study are as follows. As compared to WT ones, Mut hearts were characterized by: 1) a lower Kd_Ca_ (higher Ca^2+^ affinity) of SERCA2a ATPase activity (**Figure 1b**); 2) intracellular Ca^2+^ dynamics compatible with enhanced SR Ca^2+^ uptake (**Figure 2**); 3) insensitivity of Ca^2+^ dynamics to PLN antagonism by PST-3093 (**Figures S3, S4, and S5**); 4) lower NCX conductance (**Figure S2**). All these findings point to diminished SERCA2a inhibition by Mut PLN, resulting in increased Ca^2+^ cycling by the pump. These changes in intracellular Ca^2+^ handling were accompanied by an overall depression of energy metabolism, consisting in a parallel reduction of OCR and anaerobic glycolysis (**Figures 4 and 5**). Nonetheless, the cell energy charge, as reported by I_KATP_ conductance, was not significantly changed (**Figure S8**) and energy starvation signalling (pAMPK/AMPK) was not activated (**Figure 6**). In spite of the apparent oxidative impairment, ROS content and mitochondrial membrane polarization were remarkably normal in Mut CMs (**Figure S7**). While insensitive to cytosolic Ca^2+^ buffering, OCR was decreased by SERCA2a inhibition in WT CMs only (**Figure S6**).

### Intracellular Ca^2+^ dynamics

The mutation-associated changes in the parameters describing intracellular Ca^2+^ dynamics of intact CMs confirm the hyperdynamic state we observed in hiPS-CMs of a patient carrying the PLN-R14del mutation,^10^ thus allowing generalization of that observation. The view that hyperdynamic Ca^2+^ handling reflects enhanced SERCA2a activity is reinforced by 1) increased Ca^2+^ sensitivity (lower Kd_Ca_) of SERCA2a ATPase activity in myocardial homogenates; 2) similarity with the effect of PST-3093, a SERCA2a stimulator. The latter is known to increase SERCA2a activity by weakening its interaction with PLN,^9^ thus suggesting reduced affinity for SERCA2a as a mechanism of PLN R14del effects. In the PLN R14del heterozygous state, PST-3093 should have retained part of its stimulatory effect by displacing residual WT PLN. However, both in the PLN-R14del mouse (present study) and in heterozygous PLN R14del^+/-^ patient-derived hiPS-CMs,^10^ PST-3093 lost its SERCA2a stimulating effect completely (**Figures S3, S4, and S5**). This may suggest negative dominance of the mutation, one putative mechanism being enhanced PLN trapping in the non-inhibitory pentameric form.^21^

The hyperdynamic Ca^2+^ handling observed in the present study contrasts with the depressed Ca^2+^ handling reported in a heterozygous PLN R14del mouse based on knock-in of human PLN genes. The development of cardiac abnormalities has a rather different time-course in the two models: whereas in the former signs of depressed contractility appear at 18-20 months of age,^12^ clear-cut chamber dilatation and electrical remodelling are already present at 3 months in the latter.^22, 23^ The age at which myocyte studies were carried out is roughly similar in the two studies (2-3 months), i.e. in a markedly different relationship with the development of a failing phenotype. As depression of SR function is a landmark of the remodelled myocardium,^24^ this might explain the different Ca^2+^ handling phenotypes observed in the two models.

SERCA2a downregulation is a common consequence of maladaptive remodelling and plays an undisputable role in evolution of contractile dysfunction and arrhythmogenesis in heart failure. By analogy, enhanced SERCA2a inhibition would easily explain the PLN R14del^+/-^ ACM phenotype, as previously claimed.^3^ The present results suggest instead that PLN R14del^+/-^ loses the ability to inhibit SERCA2a; the resulting “hyperdynamic” Ca^2+^ handling may look unlikely as a cause of reduced contractility. PLN knockout, and the resulting “hyperdynamic state” of Ca^2+^ handling, is reportedly well tolerated in TG mice even with aging.^25^ However, some human mutations with PLN loss of function are associated with ACM,^26^ PLN-sarcolipin double knock-out mice undergo hypertrophic remodeling^27^ and significant derangements occur 2 months after PLN knock out in hiPS-CMs.^28^ This suggests that constitutively unrestrained SERCA2a function may, in the long term, have a negative impact on myocyte biology. Notably, clinical ACM is indeed of late onset in PLN R14del^+/-^ carriers.^1^

The discussion thus far assumes that PLN R14del detrimental effects depend on the impact of SERCA2a dysregulation on intracellular Ca^2+^ dynamics; however, this is not necessarily the case. For instance, in the case of the PLN R9C mutation, late ACM developed independently of the sign of changes in SERCA2a function.^29^ Thus, mechanisms linked to the PLN mutation, but perhaps independent of Ca^2+^ dynamics, may derange myocyte biology. Unfortunately, this considerably broadens the array of mechanisms to be considered and, eventually, to be targeted with therapy (other than mutation reversal). Hence, we undertook a preliminary analysis of additional cell dysfunctions, potentially involved in the ACM of the PLN R14del^+/-^ TG mouse. Although admittedly far from exhaustive, this analysis may provide relevant clues.

### Energy metabolism

Recent work on contracting hiPS-CMs organoids (EHTs) from a PLN R14del^+/-^ carrier^11^ detected abnormalities of the ER/mitochondrial compartment, reduced mitochondria number and function. Contraction force was halved in this preparation, in spite of a nearly normal SR function. Hence, we tested whether PLN R14del^+/-^ TG affects energy metabolism in the mouse model, notably prior to the development of contractile dysfunction (i.e. as a primary abnormality). At least when quiescent (as dictated by the experimental setup), PLN R14del^+/-^ CMs showed a substantial downregulation of energy metabolism, including its oxidative and anaerobic components (**Figures 4 and 5**).

Compatibility of overall depression of energy metabolism with grossly normal cardiac function in-vivo^12^ is more surprising. Either function is maintained despite energy starvation, or ATP demand is reduced (at least at rest) in PLN R14del^+/-^ CMs. To discriminate between these possibilities, we measured the I_KATP_ conductance (**Figure S8**), a surrogate of the ADP/ATP ratio, and activation of AMPK a major “energy starvation” signal (**Figure 6**). Albeit some heterogeneity was observed in I_KATP_ density, neither of the two measurements points to a mismatch between ATP production and demand. Why should PLN R14del^+/-^ CMs consume significantly less ATP than WT ones? Under quiescence, the condition of the metabolic measurements, the Na^+^/K^+^ pump dominates as energy consumer.^30^ Sarcolemmal Ca^2+^ extrusion, by the energetically-coupled NCX-Na^+^/K^+^ pump complex, costs twice as much ATP as Ca^2+^ recycling to the SR by SERCA2a.^31^ Therefore, lowering of SERCA2a Kd_Ca_, as in PLN R14del^+/-^ CMs, might theoretically alleviate the load on the Na^+^/K^+^ pump and be energy saving. The reduction in NCX “conductance” (**Figure S2**), possibly an adaptation to SERCA2a dominance, would contribute to limit energy consumption. This interpretation requires SERCA2a to be active at rest (see below), the condition under which OCR was measured. The hypothesis of reduced ATP demand, as opposed to impaired mitochondrial function, also fits with the presence in PLN R14del^+/-^ CMs of normal ROS levels and mitochondrial membrane polarization (**Figure S7**), both suggesting basal mitochondrial function integrity.

### Coupling between Ca^2+^ dynamics and energy metabolism

Mitochondrial Ca^2+^ plays a pivotal role in the regulation of mitochondrial respiration and ATP synthesis. To test the involvement of altered Ca^2+^ dynamics in the reduction of (resting) OCR, the latter was measured in the presence of cytosolic Ca^2+^ chelation (BAPTA), or “functional deletion” of the SR store (THAPSI). Whereas BAPTA effect was negligible in both genotypes, THAPSI significantly decreased mitochondrial basal respiration in WT only (**Figure S6**). We conclude that in WT CMs, even if quiescent, SERCA2a did contribute to OCR; furthermore, mitochondrial Ca^2+^ supply relied on a functional SR. Notably, THAPSI effect was negligible in Mut CMs, consistent with disruption of SR-mitochondrial coupling, recently described in PLN R14del^+/-^.^11^ As expected from saturation of electron transport, maximal respiration was less sensitive than basal one to THAPSI and the difference between genotypes did not achieve significance.

Comparison between BAPTA and THAPSI effects also leads to conclude that, at least under quiescence, the SR, as opposed to bulk cytosol, may be the source for mitochondrial Ca^2+^.

### Transcript and protein analysis

Cell damage might be caused, independent of SERCA2a dysregulation, by direct toxicity of the mutant protein (ER-stress) and the resulting activation of the “unfolded protein response” (UPR).^32^ ER-stress activation and reduced mitochondria abundance were indeed the main derangement detected in PLN R14del^+/-^.^11^ In the present study, transcript and protein analysis (**Figures 6** **and S9**) mainly yielded negative results, which may nonetheless be informative. Failure to detect activation of energy starvation signalling is relevant to the interpretation of depressed energy metabolism and in line with unchanged cell energy charge (normal I_KATP_ density). Lack of activation of ROS scavenging genes, along with normal ROS content and mitochondrial membrane potential, stand for the absence of major function alteration under basal condition. Similarly, lack of CaMKII activation argues against early involvement of Ca^2+^ decompartmentalization in disease pathogenesis.

## 5. LIMITATIONS

Because of technical constraints, energy metabolism was measured in quiescent CMs. Although the response of OCR to Thapsigargin confirms partial activation of SERCA2a even in this condition, the cell metabolic state might be entirely different in contracting myocytes. Accordingly, the results of metabolic measurements should not be taken to rule out energy deficiency as a pathogenetic mechanism in contracting hearts. They may rather reflect deficient regulation of mitochondrial function by Ca^2+^ in mutant myocytes.

## 6. CONCLUSIONS

The present results extend our previous observation in patient-derived hiPS-CMs^10^ to indicate that PLN R14del^+/-^ may upregulate SERCA2a activity and induce hyperdynamic Ca^2+^ handling in native mature CMs of the TG mouse. The mutation also results in a substantial downregulation of energetic metabolism, not associated with signs of mitochondrial dysfunction or energy decompensation. The simplest comprehensive interpretation of these findings would see downregulation of energetic metabolism in mutant CMs as the “physiological” response to reduced ATP demand, expected from switch to SERCA2a dominance in Ca^2+^ handling. Nonetheless OCR insensitivity to SERCA2a blockade (THAPSI) suggests impairment of Ca^2+^ transfer from SR to mitochondria as a further cause of OCR downregulation in mutant CMs. At any rate, albeit with the limitation inherent to measurements in quiescent CMs (see above), metabolic incompetence seems unlikely as a primary pathogenetic mechanism.

Albeit reportedly well tolerated in mice,^25^ hyperdynamic Ca^2+^ handling is nonetheless abnormal. Notably, the metabolic response to PLN knock out in hiPS-CMs is a transient increase in OCR, with mitochondrial damage appearing only as a late consequence.^28^ Both PLN R14del^+/-^ and PLN knock out induce a hyperdynamic state, but in the latter PLN protein is missing. This may suggest an additional pathogenetic mechanism, independent from changes in SERCA2a function: ACM might result from toxicity of the mutant protein. Indeed, intracellular PLN deposits are a prominent feature in overt ACM, clear-cut ER stress has been reported in PLN R14del^+/-^ hiPS-CMs^11^ and, in spite of the early disease stage, the present findings also suggest activation of unfolded protein response (UPR). Once instated, UPR can be responsible for a host of cellular abnormalities, including major mitochondrial remodelling and SR damage.^32^ Since many of these processes are Ca^2+^-dependent, the view that ACM pathogenesis might be independent of Ca^2+^ handling dysregulation may not conflict with the observation that strong Ca^2+^ buffering may prevent phenotype development.^11^

Last but not least, the cellular mutation phenotype of the TG mouse described in the present study is, under most aspects, consistent with that of patient-derived hiPS-CMs,^10, 11^ thus providing cross-validation of these experimental models.

## 7. THERAPEUTIC IMPLICATIONS

The initial hypothesis of SERCA2a superinhibition by PLN R14del^+/-^ pointed to the therapeutic potential of PLN-displacing SERCA2a activators, now available as drugs.^9^ ^18^ ^33^ The loss of SERCA2a function detected in the zebrafish TG model of the mutation, was indeed reversed by one of these compounds (istaroxime).^7^ However, consistent with a loss of SERCA2a inhibition by PLN R14del^+/-^, the prototypical selective SERCA2a activator PST-3093, was totally ineffective in patient-derived hiPS-CMs^10^ and TG mouse CMs. This, and the likely involvement of pathogenetic mechanisms beyond SERCA2a dysregulation, may suggest to invest on mutation reversal,^34^ or at least on improvement of mutant protein processing (e.g. by “chaperone” molecules), as the most logical mechanism-based therapeutic approaches. Speaking of more generic approaches, countering the enhancement of sustained Na^+^ current (a common response to cell stress) has proven effective in preventing the consequences of chronic upregulation of Ca^2+^ cycling in hiPS-CMs.^28^

## 8 DECLARATIONS

## Acknowledgements

Prof. Bianchi (Windtree Therapeutics) for supplying PST3093.

This work was funded by grants from CVie Therapeutics Limited for the development of PLN antagonists (to AZ) and from the Europen Union-Italian Ministry of University and Research – grant PNRR – M4C2-I1.3 Project PE_00000019 “HEAL ITALIA” (to ES). The Operetta and Seahorse platforms were acquired through the MUR-Competitive Grant for Excellent Departments (2018-2022) to University Milano-Bicocca. The PLN-R14del murine model was funded by grants of the Netherlands Heart Foundation (CVON PREDICT2, grant 2018-30), the leDucq Foundation (Cure PhosphoLambaN induced Cardiomyopathy (Cure-PLaN), and the NLHI and the Dutch PLN Foundation to RdB and HS.

## Conflicts of interest / Declarations

AZ recipient of research funding by CVie Therapeutics Limited (Taipei, Taiwan), WindTree Therapeutics (Warrington, USA) for research on PLN antagonists.

The UMCG which employed/employs several of the authors received research grants and/or fees from AstraZeneca, Abbott, Boehringer Ingelheim, Cardior Pharmaceuticals GmbH, Ionis Pharmaceuticals, Inc., Novo Nordisk, and Roche. Dr. de Boer has had speaker engagements with Abbott, AstraZeneca, Bayer, Bristol Myers Squibb, Novartis, and Roche.

The views and opinions expressed in this manuscript are those of the authors only and do not necessarily reflect those of the European Union or the European Commission. Neither the European Union nor the European Commission can be held responsible for them.

## Availability of data and material

The authors confirm that the data supporting the findings of this study are available within the article and/or its supplementary materials. The data that support the findings of this study are available from the corresponding author upon request.

