## Supplementary Material for "Early consequences of the phospholamban mutation PLN-R14del^+/-^ in a transgenic mouse model"

**SUPPLEMENT**

**DETAILED MATERIALS AND METHODS**

***Experimental model***

Development of the PLN R14del TG mouse is described by Eijgenraam et al.^1^ The studies were performed on tissues from PLN R14del heterozygous (Mut) mice and WT controls aged 8 to 12 weeks. In this age range, the animals are healthy; overt physical or echocardiographic signs of chamber remodelling or contractile dysfunction are present in this model at 18 months of age.^1^ This is crucial to the interpretation of the observed changes as directly resulting from the mutation, rather than secondary to aspecific myocardial remodelling.

***Cardiomyocyte isolation***

All experiments involving animals confirmed to the guidelines for Animal Care endorsed by the Milano-Bicocca and to the Directive 2010/63/EU of the European Parliament on the protection of animals used for scientific purposes.

Mut mice and WT controls were intraperitoneally anesthetized with chloralhydrat (ClCCH(OH)_2_, Sigma - Aldrich) in 200 μL of Seleparina (Nadroparina Calcica 9500 U.I, ItalFarmaco) and 200 μL of physiological solution (0.9% NaCl 1M in MilliQ). After anaesthesia, evidenced by absence of movements and reflexes, animals were sacrificed by cervical dislocation and ventricular cardiomyocytes (CMs) were isolated by using a manual perfusion method.^2^

Briefly, while the heart was still in situ, the right ventricle (RV) was injected with 7 mL of EDTA buffer solution (130 mM NaCl, 5 mM KCl, 0.5 mM NaH_2_PO_4_, 10 mM HEPES, 10 mM Glucose, 10 mM 2,3-Butanedione Monossime, BDM, 10 mM Taurine, 5 mM EDTA; pH=7.8 with NaOH); the ascending aorta was then clamped and the heart was transferred to a dish containing EDTA buffer solution.

Digestion was achieved by sequential injections into the left ventricle (LV) of 10 mL EDTA buffer solution, 6 mL of perfusion buffer solution (130 mM NaCl, 5 mM KCl, 0.5 mM NaH_2_PO_4_, 10 mM HEPES, 10 mM Glucose, 10 mM 2,3-Butanedione Monossime, BDM, 10 mM Taurine, 1 mM MgCl_2_; pH=7.8 with NaOH) and 20 to 30 mL of Liberase buffer solution (0.1 mg/ml Liberase in Perfusion Buffer + 0.02 mg/ml Trypsin EDTA and 6.25 µM CaCl_2_).

Cellular dissociation was completed by gentle trituration, while enzyme activity was quenched by adding 10 mL of stop buffer solution (perfusion Buffer + 10% FBS). The resulting suspension was filtered, centrifuged and resuspended in calcium free solution (130 mM NaCl, 5.4 mM KCl, 0.4 mM NaH_2_PO_4_, 0.5 mM MgCl_2_, 25 mM Hepes, 22 mM Glucose; pH=7.4 with NaOH). Finally, the physiological extracellular Ca^2+^ concentration (2 mM) was gradually reestablished by adding to the cell suspension small volumes at increasing CaCl_2_ concentration.

***Preparation of cardiac homogenate***

For each group of tissues, the hearts were divided into two pools to be processed separately, in order to get two different preparations.

The hearts were thawed, weighed, dissected by surgical scissor, and homogenized 4 times (20 seconds for each period) with IKA UltraTurrax at 4°C in homogenization buffer (ratio 1g / 4 ml) containing 50 mM Potassium Phosphate, 300 mM Sucrose, Sodium Fluoride 10 mM, pH 7 plus 0.3 mM PMSF and 0.5 mM DTT. The total homogenate was centrifuged at 17000 rpm for 30 minutes at 4°C (Beckman); the final pellet was resuspended in 2.5 volumes of homogenization buffer.

***SERCA2a ATPase measurements in cardiac homogenates***

The measurement was performed as previously described.^3^ Briefly, SERCA2a ATPase activity was measured in homogenates as the rate of ^32^P-ATP hydrolysis and identified as the fraction inhibited by 10 μM cyclopiazonic acid (CPA). The ATPase activity was measured at multiple Ca^2+^ concentrations; data points were fitted to a sigmoidal (Hill) function, from which the maximum velocity (V_max_) and Ca^2+^ affinity [Kd_Ca_] parameters were estimated.

***Electrophysiological recordings***

*Ruptured-patch recordings*

Except for I_KATP_ measurements, V-clamp was performed by the ruptured-patch version of the whole-cell mode. Cells were perfused (at 36 °C) with an extracellular solution containing (in mM): 154 NaCl, 4 KCl, 5 HEPES NaOH, 1 mM CaCl_2_, 1 mM MgCl_2_, 5.5 glucose; pH=7.35 with NaOH. Patch pipettes (tip resistance of 1.5-2.5 MΩ) were filled with an intracellular solution containing (in mM): 110 K-aspartate, 23 KCl, 3 MgCl_2_, 0.04 CaCl_2_, 0.1 EGTA KOH (10^-7^ Ca^2+^-free), 5 Hepes KOH, 0.4 Na^+^-GTP, 5 Na^+^-ATP, 5 Na^+^-phosphocreatine; pH=7.3 with KOH. Membrane capacitance and series resistance (<10 MΩ) were measured in every cell but left uncompensated. The V protocols were applied, and the recorded signals amplified by MultiClamp 700B (Molecular Devices, Sunnyvale, CA, United States), digitized at 20 kHz (Axon Digidata 1440A, Molecular Devices), and filtered at 10 kHz. As validated in previous work, membrane current (I_m_) recorded during steps from -40 to 0 mV in the presence of 2 mM 4-aminopyridine and 0.1 mM BaCl_2_ are largely representative of I_CaL_; intracellular K^+^ was not substituted to avoid interference with SR function. The change in I_m_ induced by caffeine challenge (at V_m_ = -40 mV) was taken as a measure of Na^+^/Ca^2+^ exchanger current (I_NCX_).^4^

*Perforated-patch recordings*

I_KATP_ was recorded using amphotericin-perforated patch technique, aimed to prevent intracellular contamination by the pipette solution.^5^ Briefly, a stock solution was prepared by sonicating 3 mg amphotericin B (Merck) in 50 μl dimethylsulphoxide (DMSO). Prior to each experiment, 3.33 μl amphotericin stock was added to 1 ml of intracellular solution. To enhance seal formation, the pipette tip was dipped in amphotericin-free intracellular solution for 20 s prior to backfilling with the amphotericin containing one. Following seal formation, small depolarizing pulses (10 mV from -80 mV) were applied and overall “access resistance” was computed from the capacitive current transients. The latter became stable after about 10 min; poration was considered satisfactory when access resistances fell below 25 MΩ.

I_KATP_ was evaluated at a holding potential of -120 mV as the difference (I_CTRL_ – I_GLIBE_) between the current recorded in control (CTRL) and during steady-state perfusion with the selective blocker glibenclamide (GLIBE, 5 μM). I_KATP_ was normalized to membrane capacitance to obtain current density.

***Intracellular Ca^2+^ recordings***

Intracellular Ca^2+^ measurements were performed at 36°C. Cytosolic Ca^2+^ was optically measured in ventricular CMs loaded with the probe Fluo-4 AM (10 µM) by 30 min incubation at room temperature. During the recording, cells were perfused with an extracellular solution containing (in mM): 154 NaCl, 4 KCl, 5 HEPES NaOH, 2 mM CaCl_2_, 1 mM MgCl_2_, 5.5 glucose; pH=7.35 with NaOH.

Calcium transients (CaT) were recorded as Fluo-4 signal, collected at 40X magnification (oil immersion) through a 535 nm band pass filter, converted to voltage, low-pass filtered (10 kHz) and digitized at 20 kHz. To compensate for potential differences in probe loading, all fluorescence (F) values were normalized to F0, which was recorded during prolonged quiescence.

Ca^2+^ transients (CaT) were measured either under field-stimulation (1 Hz) or V-clamp, as specified in the results section. The following parameters were extracted from CaT: Ca^2+^ transient amplitude (CaT amplitude), Ca^2+^ transient decay kinetics (τ _decay_), Ca^2+^ transient rise-time (t_peak_), diastolic Ca^2+^ (CaD). Rate-dependency of CaT properties was tested by stepwise increments in pacing rate (to 1, 1.3, 1.7, and 2 Hz). Under V-clamp, simultaneous measurement of I_CaL_ allowed to estimate the “gain” of Ca^2+^-induced Ca^2+^ release (CICR) as the Ca^2+^ release/influx ratio.^6^

The sarcoplasmic reticulum (SR) Ca^2+^ content was estimated as the integral of the I_m_ (mostly representing I_NCX_ ) elicited by a caffeine (10 mM) pulse applied after a loading train of V steps (−40 to 0 mV at 1 Hz).^4^ To avoid Ca^2+^ influx during the SR emptying, caffeine was dissolved in Ca^2+^-free solution (containing 1 mM EGTA CsOH).

The fraction of SR Ca^2+^ content released by membrane excitation (fractional release) was calculated as the ratio between the amplitudes of V- and caffeine-triggered CaT. The “gain” of Ca^2+^ -induced Ca^2+^ release (CICR) was measured as the ratio between Ca^2+^ influx through I_CaL_ and the resulting increase in cytosolic Ca^2+^.

Information on NCX function was obtained by linear fitting of the trailing branch of the I_NCX_/[Ca^2+^] loops, recorded during caffeine-induced transients. The slope coefficient and the 0 I_NCX_ intercept were used as surrogate of NCX “conductance” and cytosolic [Ca^2+^] at electrochemical equilibrium, respectively.

The Ca^2+^ uptake function of the SR was evaluated through a dedicated “SR reloading” protocol, applied under V-clamp.^7^ The SR was initially emptied by a caffeine pulse and reloaded by a train (0.25 Hz) of 200 ms V steps from −40 to 0 mV. CaT and I_CaL_ were simultaneously recorded during the reloading protocol. The following parameters were analysed from each step of the protocol: i) CaT amplitude, ii) CICR gain, iii) CaT decay kinetics (τ _decay_), iv) diastolic Ca^2+^ (CaD). The rate of increment of the former two parameters during the loading protocol reports the rate of SR refilling. τ _decay_ reports the rate of cytosolic Ca^2+^ clearance (the faster Ca^2+^ removal, the smaller τ _decay_) within each step, i.e. at varying SR filling levels. CaD course reports the rate of cytosolic Ca^2+^ accumulation, expectedly reduced by enhancement of SR uptake function.

***Analysis of mitochondrial respiration and anaerobic glycolysis***

Oxygen consumption rate (OCR; an index of oxidative phosphorylation) and Proton Efflux Rate (PER; an index of anaerobic glycolysis) were measured within each plate in quiescent CMs (Seahorse XFe96 Extracellular Flux Analyzer, Agilent) by using the Mito Stress Test and the Glycolytic Rate Assay protocols, respectively. To minimize measurement stray variability, both experimental groups were represented within the same multiwell plate; furthermore, the position of each treatment within the plate was changed in each session. To prevent cell displacement, plate wells were coated with the adhesive agent CellTak (22.4 μg/ml, Corning). Cell concentration in the CMs suspension was preliminarily analysed (Operetta CLS™– Perkin Elmer). Viable cells per unit volume were counted as the difference between number of nuclei (stained by Hoechst 0.5 μg/ml) and of dead cells stained with Propidium Iodide (PI 0.5 μg/ml). Based on the count, CMs suspension (DMEM Medium, Agilent Seahorse XF, pH=7.4) was seeded into each well to obtain a density of 2.5×10^3^ cells/well.

OCR was measured under basal conditions and after injection of oligomycin (Oligo; 1.5, 3 μM), carbonyl cyanide *p*-trifluoromethoxyphenylhydrazone (FCCP; 0.5, 1, 2 μM) and Rotenone/Antimycin A (Rot/Ant; 0.5, 1, 3 μM). The minimal doses of drugs causing the maximal response were determined for each experimental group in preliminary measurements.

OCR parameters were calculated as follows: i) mitochondrial basal respiration = OCR before Oligo (1.5 μM) – OCR after Rot/Ant (3 µM); ii) mitochondrial maximal respiration = OCR after FCCP (0.5 µM) − OCR after Rot/Ant (3 µM); iii) spare respiratory capacity = mitochondrial maximal respiration – mitochondrial basal respiration; iv) non-mitochondrial respiration = OCR after Rot/Ant (3 µM).

PER was measured under basal conditions and after injection of 1.5 μM Oligo, 3 μM Rot/Ant and 50 mM 2-deoxy-d-glucose (2-DG) to assess anaerobic glycolysis.

Glycolytic parameters were calculated as follows: i) basal glycolysis = PER before Oligo – PER after 2DG); ii) compensatory glycolysis = PER after Rot/Ant − PER after 2-DG; iii) glycolytic reserve = compensatory glycolysis – basal glycolysis; iv) non-glycolytic acidification = PER after 2-DG.

At the end of the experiment, measured OCR and PER values were normalized to the number of viable cells in each well, again quantified as above.

To assess the influence on OCR of cytosolic and SR Ca^2+^ respectively, a subset of cells was pre-incubated for 30 minutes with BAPTA-AM (BAPTA; 5 μM) or Thapsigargin (THAPSI; 5 μM). Vehicle (DMSO) was present at the same concentration in all the experimental solutions.

***ROS quantification***

To evaluate intracellular content of reactive oxygen species (ROS), quiescent CMs were stained with 2’7’-dichclorofluorescin diacetate (DCFDA; 20 μM) for 10 minutes at 37°C and 5% CO_2_. Stained cells were seeded, at the concentration of 2.5×10^3^ per well, in 96-wells plates (Perkin Elmer). DCFDA was excited at 485 nm and ROS levels were estimated as fluorescence emission at 535 nm. Automated (unbiased) confocal single-cell fluorescence measurement was performed by Operetta CLS™ (Operetta– Perkin Elmer) at 40× magnification.

***Mitochondrial Membrane Potential evaluation***

To evaluate mitochondrial membrane potential (Ψ_m_), cells were stained with 100 nM Tetramethylrhodamine, Ethyl Ester (TMRE) for 10 min at 37 °C and 5% CO_2_. Stained cells were seeded at the concentration 7.5×10^3^ per well. TMRE fluorescence was analysed by confocal imaging at 63× magnification (Operetta CLS™). At the concentration used in the present study, TMRE works in “quenching mode”, i.e. emission increases as Ψ_m_ depolarizes.^8^ At any rate, TMRE fluorescence signal was calibrated in each measurement by short-circuiting mitochondrial electron transport with FCCP (0.02 μM).

***mRNA extraction and qRT-PCR assay***

After collection, RV bioptic samples from WT and Mut mice were mechanically disrupted using metallic-beads by a TissueLyser (Qiagen, Milan, Italy) in an appropriate amount of RL lysis buffer (Norgen Biotek corp., Thorold, Canada). RNA was extracted by using a total RNA purification kit (Norgen Biotek corp., Thorold, Canada) and quantified by a NanoDrop spectrophotometer (ND-1000, EuroClone, Milan, Italy). Reverse transcription was conducted with SuperScript III (Invitrogen, Carlsbad, CA, USA), following manufacturer’s instructions. qRT-PCR was performed with the iQTM SYBR Green Super Mix (Bio-Rad Laboratories, Hercules, CA, USA) reagent and specific primers (reported in **Table 1**). All reactions were performed in a 384-well format with the 7900HT Fast Real-Time PCR System (Thermo Fisher Scientific, Massachusetts, USA). The relative quantities of specific mRNAs were obtained by the comparative Ct method and normalized to the housekeeping gene glyceraldehyde 3-phosphate dehydrogenase (GAPDH).

***2.11 Protein Extraction and Western Blot Analysis***

RV samples were chopped using metallic-beads by a TissueLyser (Qiagen, Milan, Italy) in cell lysis buffer (Cell Signaling Technology, Danvers, MA, USA) supplemented with protease- and phosphatase-inhibitor cocktails (Sigma-Aldrich, Saint Louis, MO, USA). Protein lysate was then quantified, and equal amounts of total protein extracts were subjected to SDS-PAGE and transferred onto a nitrocellulose membrane (Bio-Rad, California, USA). The membranes were blocked for 1 hour at room temperature in 5% non-fat dry milk added to Wash Buffer (Tris Buffer Sulfate, 0.1% Tween-20) and then incubated overnight at 4°C with the appropriate primary antibodies (reported in **Table 2**). Peroxidase-conjugated secondary antibodies (GE Healthcare, Chicago, IL, USA) were then applied for 1 hour. Peroxidase signal was visualized using the LiteUP Western Blot Chemiluminescent Substrate (EuroClone, Milan, Italy). Images were acquired with the ChemiDoc^TM^ MP Imaging System (Bio-Rad, California, USA), and blot densitometric analysis was performed (ImageJ software, National Institutes of Health, Bethesda, MD, USA). Protein signals were normalized to GAPDH and TUBULIN based on the gel gradient used.

***2.12 Statistical analysis***

GraphPad Prism 8 (GraphPad software, San Diego, CA, USA) was used for statistical analysis.

Normality of distribution was assessed using D’Agostino-Pearson’s normality test. To compare two sample means, either the Student’s t-test and the Mann-Whitney U-test were used for continuous or categorical data, respectively. To compare more than two sample means, one-way ANOVA (with Tukey correction) or Kruskal-Wallis (with Dunn’s correction) were used for continuous or categorical data, respectively. In the case of repeated measurements (e.g. rate-dependency and SR loading) a Mixed-effects model containing “Treatment” (WT vs Mut, PST-3093 vs Control) and “Variable” (rate or step #) factors was used. As reported for each result, the significance of “Treatment X Variable interaction” (i.e. difference between Treatments in their response to the Variable) was first tested; in its absence, significance of difference between Treatments at all Variable values was tested.

In figures, whenever feasible, individual data points were plotted, to illustrate dispersion, along with the sample mean ± SEM. Whenever the threshold for statistical significance (p<0.05) was achieved, the actual p-value for the comparison was reported as an index of robustness. For each experiment, the number of preparations or cells (n) and the number of animals from which they were obtained (N) are indicated in the respective figure legend.

**TABLES**

***Table 1. Primer sequences 5’ - 3’.***

| **Gene** | **Forward primer** | **Reverse primer** |
| --- | --- | --- |
| *Eif2ak3* | CACGCAGATCACAGTCAGG | GTGGGGCTGAGGATGGAAAA |
| *Hspa5* | GGGTCAGGGAGAGGAGGAAT | CCAAGGTGAACACACACCCT |
| *Atf6* | TGGAAGCCTAAAGAGGACCTG | CGTGGGAGGACAGAGAAACAA |
| *Ern1* | GAGACAGATTGTCAGGGCCA | CCTACAAAGTCTGTGGTAGCCT |
| *Pink1* | GATGGCTCTGTGCTCCAGTT | CCCTCCCTCTACTCCAGCTT |
| *Parkin* | GGGATTCAGAAGCAGCCAGA | GAGGGTTGCTTGTTTGCAGG |
| *Map1lc3b* | CCAAGCCTTCTTCCTCCTGG | TTGCTGTCCCGAATGTCTCC |
| *Vdac1* | CGGCCACACATGATCACAGA | ACCAGTCTCGGGGTCTTCTT |
| *Itp3r 1* | GTTGGACGAGGCTGGAAATG | GGCCAGCATTGACAGGATTC |
| *Itp3r 2* | GGGAGAAATCGTGAAATACAGCA | CATCCAGTGACACTCGCATG |
| *Itp3r 3* | CTCTGTCAACTGCAACACCA | TTGCCCCTGTACTCATCACA |
| *Hspa9* | AGGTTTCCAGAAGCGTAGCC | GGTTACGAGGGCAAGAACCA |
| *Mfn2* | GCTTGGACAGGTGGAGTCAA | GGACTCGAGGTCTCCTCTGT |
| *Gpx3* | GTGAACGGGGAGAAAGAGCA | CAGGAGTTCTGCAGTGGGAG |
| *Sod2* | GTAGGGCCTGTCCGATGATG | CGCTACTGAGAAAGGTGCCA |
| *Nqo1* | GCTCCACGAGGATGGGAAAA | TGCCCTGAGGCTCCTAATCT |
| *Ho1* | CCTCACAGATGGCGTCACTT | TGGGGGCCAGTATTGCATTT |
| *Gclc* | GCTTTGGGTCGCAAGTAGGA | GCGTCCCGTCCGTTCC |
| *Ptgs2* | CTTCGGGAGCACAACAGAGT | AAGTGGTAACCGCTCAGGTG |
| *Camk2a* | ATCGTCCACTTCCACAGA | CCACATTCCACGGACAAA |
| *Camk2b* | GCATTCCAACATTGTACGC | CTTGCCACGATGTCTTCA |
| *Camk2g* | TGATGCCAGCCACTGTA | GCAAGTTCTCAGGCTTCA |
| *Camk2d* | TGTTCACTTTCACCGTTCG | GCCTTGAGGACAGAATGAAG |
| *Prkaa1* | AAAGTGAAGGTGGGCAAGCA | CAGATGGTGTACTGATGACCTGG |
| *Prkaa2* | TCGCAGTTTAGATGTTGTTGGA | CTTCAACCCGCCCATGTTTG |
| *Prkab1* | CCTTGTGTCCCTGCAGATTCC | CATGAGGATCTTGGGCCTGTC |
| *Prkab2* | CGACCCCAGCGTCTTCAG | CAAATCCTGCTGCCAGGGTA |
| *Prkag1* | GCTTTCCAAGCTGAGGAACT | ACCAACTTGGAACTTGTGGG |
| *Prkag2* | TGCCTACTTAAGGACAGGCG | CCCTGAGACTCCTGAGAAAGC |
| *Prkag3* | CCATGGCTACCAGCTCCAAA | CTCTGCTTCTTGCTGTCCCA |
| *Gapdh* | ACCACAGTCCATGCCATCAC | TCCACCACCCTGTTGCTGTA |

***Table 2. Primary antibodies.***

| **Protein** | **Clonality/Code** | **Source** | **Company** | **Diluition** |
| --- | --- | --- | --- | --- |
| phospho-CaMKII (T286) | Polyclonal, ab32678 | Rabbit | Abcam | WB: 1:1000 |
| CaMKII | Monoclonal, ab52476 | Rabbit | Abcam | WB: 1:1000 |
| phospho-AMPK | Monoclonal, #2535 | Rabbit | Cell Signaling | WB: 1:1000 |
| AMPK | Polyclonal, #2532 | Rabbit | Cell Signaling | WB: 1:1000 |
| GAPDH | Polyclonal, sc-25778 | Rabbit | Santa Cruz | WB: 1:1000 |

**SUPPLEMENTARY FIGURES**

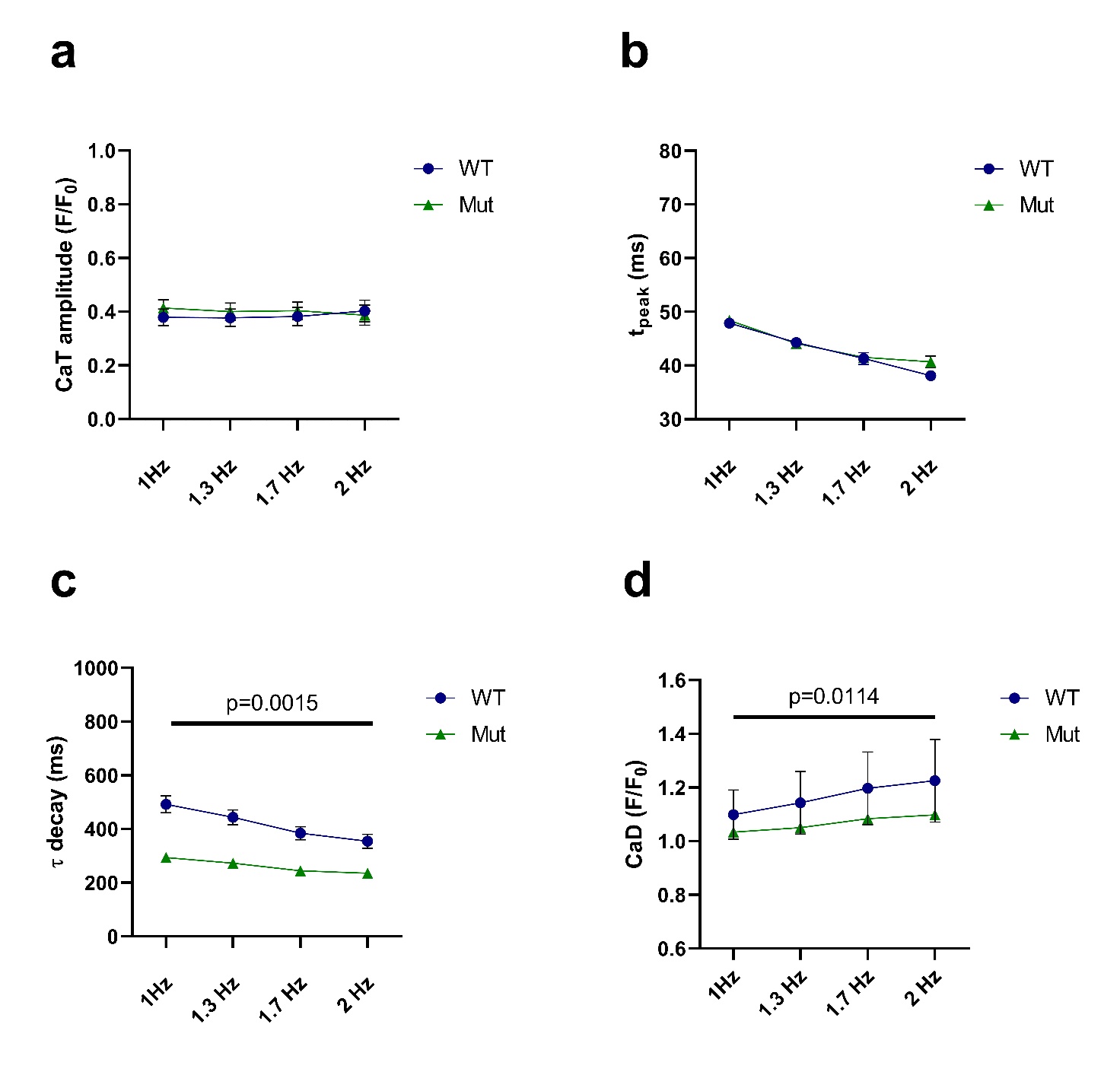

**Figure S1. Rate dependency of CaT parameters in WT and Mut.** Rate-dependency was tested by stepwise increments in pacing rate (1, 1.3, 1.7, and 2 Hz). (**a**) Ca^2+^ transient amplitude. (**b**) CaT time to peak, t_peak_. (**c**) Ca^2+^ transient decay kinetics, τ decay. (**d**) Diastolic Ca^2+^, CaD. WT: n=25, N=4; Mut: n=17, N=3. Data are expressed as mean ± SEM. Mixed-effects model, Treatment X Rate.

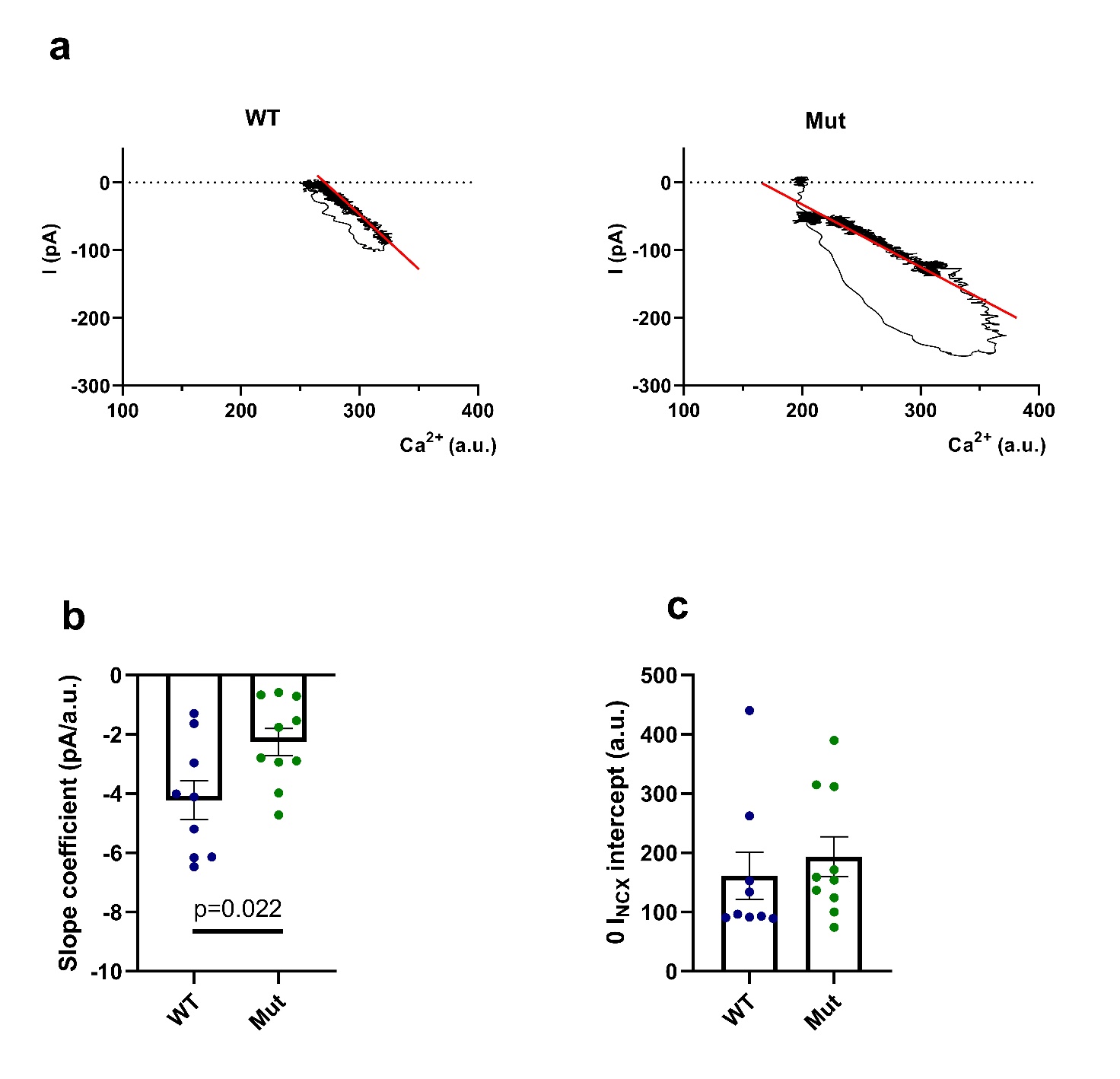

**Figure S2. Relationship between I_NCX_ and [Ca^2+^] in WT and Mut.** (**a**) Representative I_NCX_/[Ca^2+^] loops during caffeine pulse. A linear regression (red solid line) has been drawn through the trailing branch of the loop. (**b**) Line slope coefficient used to define NCX “conductance” (WT: -4.2 ± 0.65, n=9; Mut: -2.3 ± 0.46, n=10). (**c**) 0 I_NCX_ intercept indicating cytosolic [Ca^2+^] at electrochemical equilibrium (WT: 161 ± 40, n=9; Mut: 194 ± 34, n=10). WT: N=7; Mut: N=6. Data are expressed as mean ± SEM. Mann-Whitney U-test.

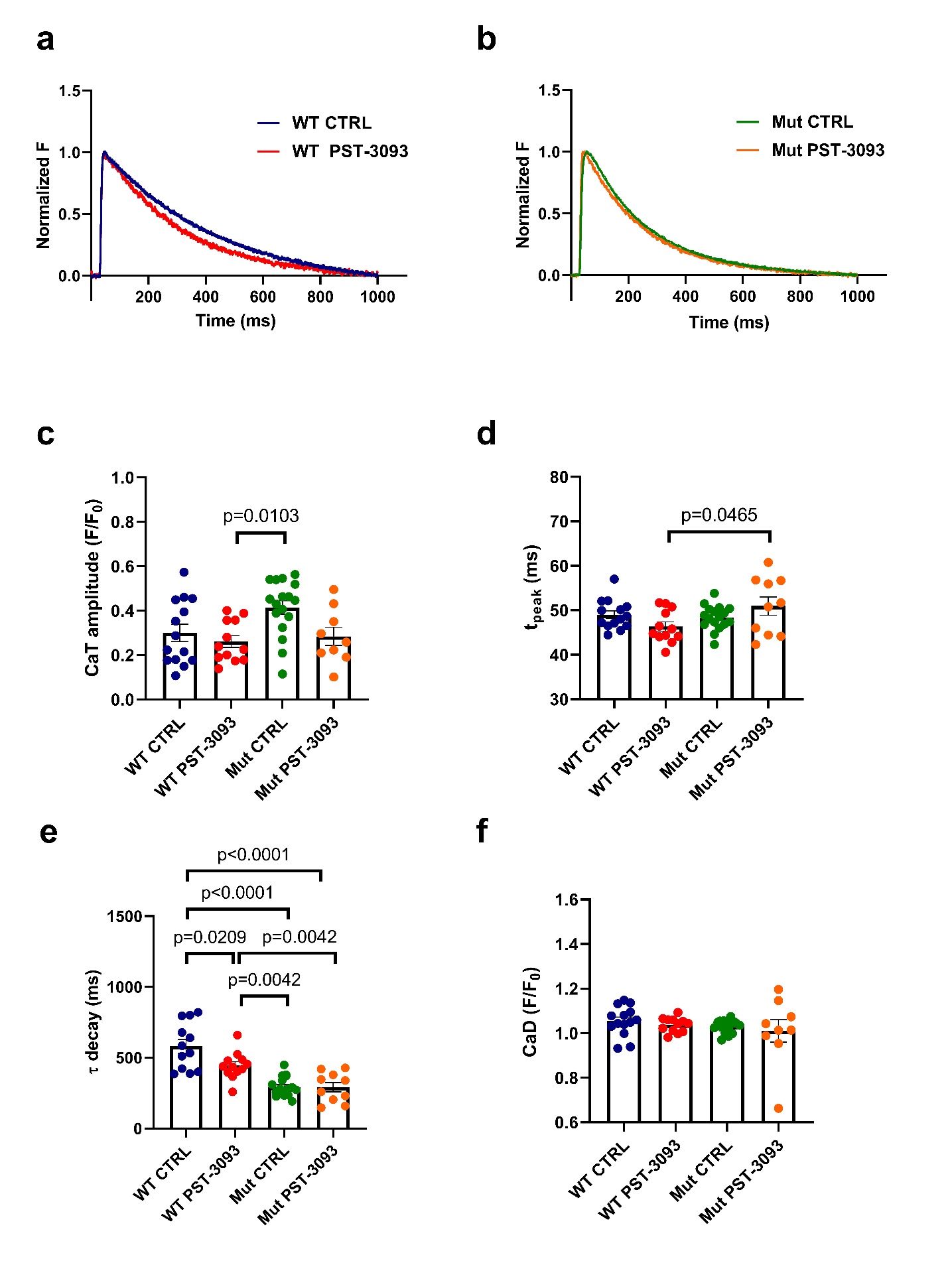

**Figure S3. PST-3093 (1µM) effect on CaT parameters in WT and Mut.** (**a-b**) Representative Ca^2+^ transient traces. (**c**) Ca^2+^ transient amplitude (WT CTRL: 0.30 ± 0.04, n=14; WT PST-3093: 0.26 ± 0.03, n=12; Mut CTRL: 0.41 ± 0.03, n=17; Mut PST-3093: 0.28 ± 0.04, n=9). (**d**) CaT time to peak, t_peak_ (WT CTRL: 49 ± 0.89, n=14; WT PST-3093: 46.4 ± 1.05, n=12; Mut CTRL: 48.4 ± 0.66, n=17; Mut PST-3093: 50.9 ± 2.05, n =10). (**e**) Ca^2+^ transient decay kinetics, τ_decay_ (WT CTRL: 580 ± 48, n=12; WT PST-3093: 444 ± 27.7, n=12; Mut CTRL: 293 ± 16.5, n=17; Mut PST-3093: 291 ± 33.1, n=10). (**f**) Diastolic Ca^2+^, CaD (WT CTRL: 1.05 ± 0.02, n=14; WT PST-3093: 1.04 ± 0.01, n=12; Mut CTRL: 1.03 ± 0.01, n=17; Mut PST-3093: 1.01 ± 0.05, n=9). WT: N=3; WT PST-3093: N=3; Mut: N=3; Mut PST-3093: N=4. Data are expressed as mean ± SEM. One-way ANOVA.

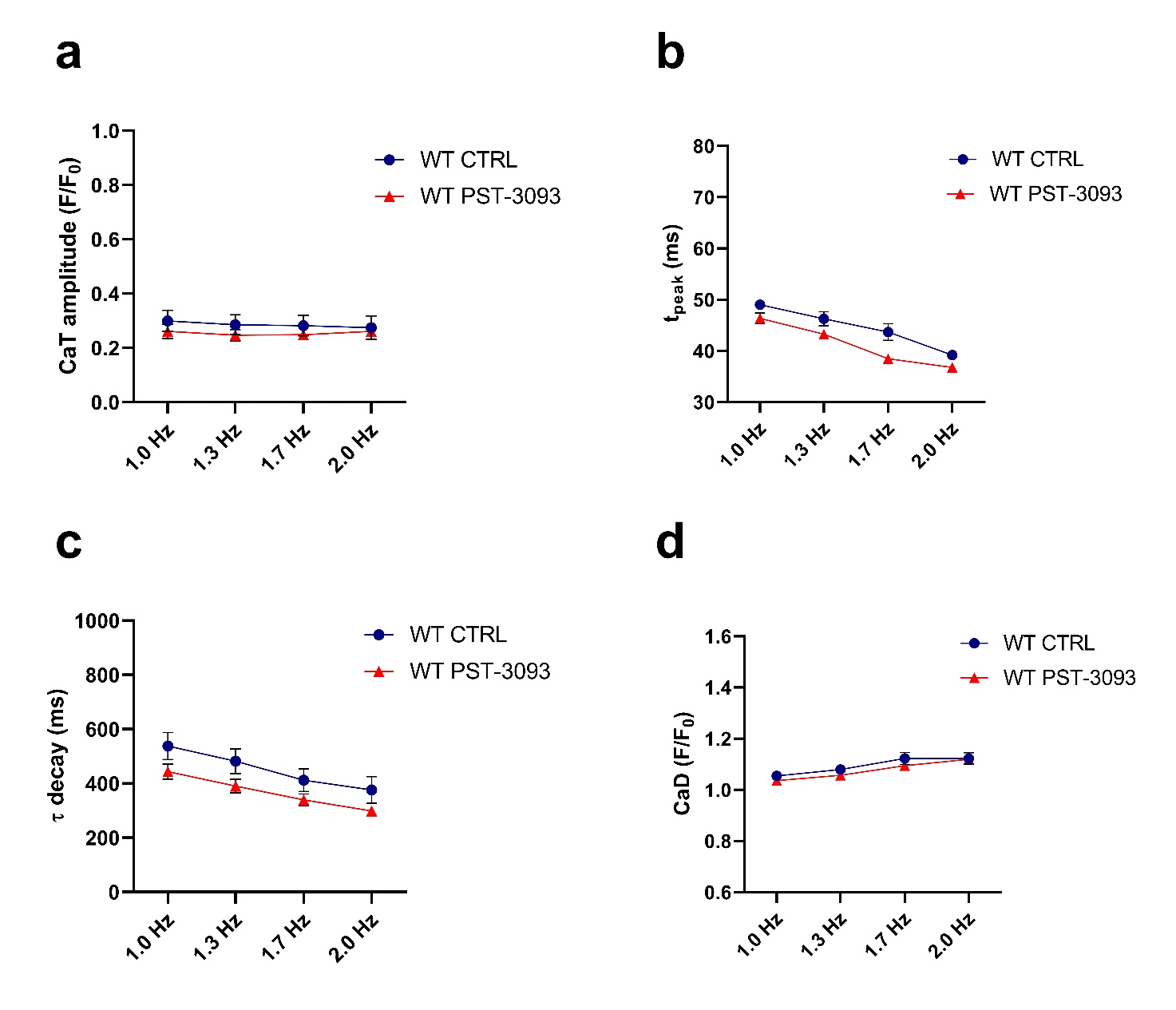

**Figure S4. PST-3093 1µM effect on rate dependency of CaT parameters in WT.** Rate-dependency was tested by stepwise increments in pacing rate (1, 1.3, 1.7, and 2 Hz). (**a**) Ca^2+^ transient amplitude. (**b**) CaT time to peak, t_peak_. (**c**) Ca^2+^ transient decay kinetics, τ decay. (**d**) Diastolic Ca^2+^, CaD. WT CTRL: n=14, N=3; WT PST-3093: n=12, N=2. Data are expressed as mean ± SEM. Mixed-effects model.

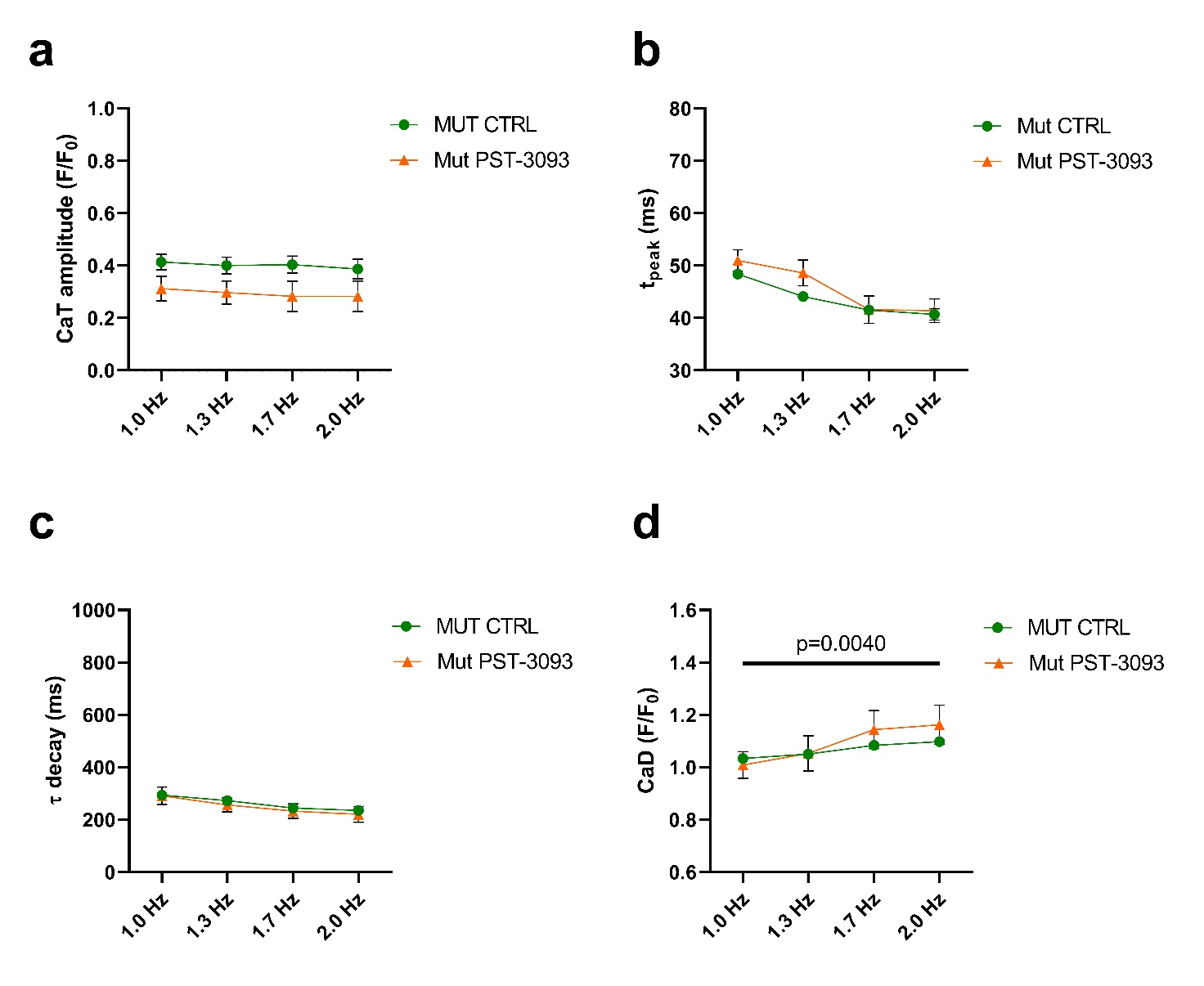

**Figure S5. PST-3093 1µM effect on rate dependency of CaT parameters in Mut.** Rate-dependency was tested by stepwise increments in pacing rate (1, 1.3, 1.7, and 2 Hz). (**a**) Ca^2+^ transient amplitude. (**b**) CaT time to peak, t_peak_. (**c**) Ca^2+^ transient decay kinetics, τ decay. (**d**) Diastolic Ca^2+^, CaD. Mut CTRL: n=17, N=3; Mut PST-3093: n=10, N=4. Data are expressed as mean ± SEM. Mixed-effects model.

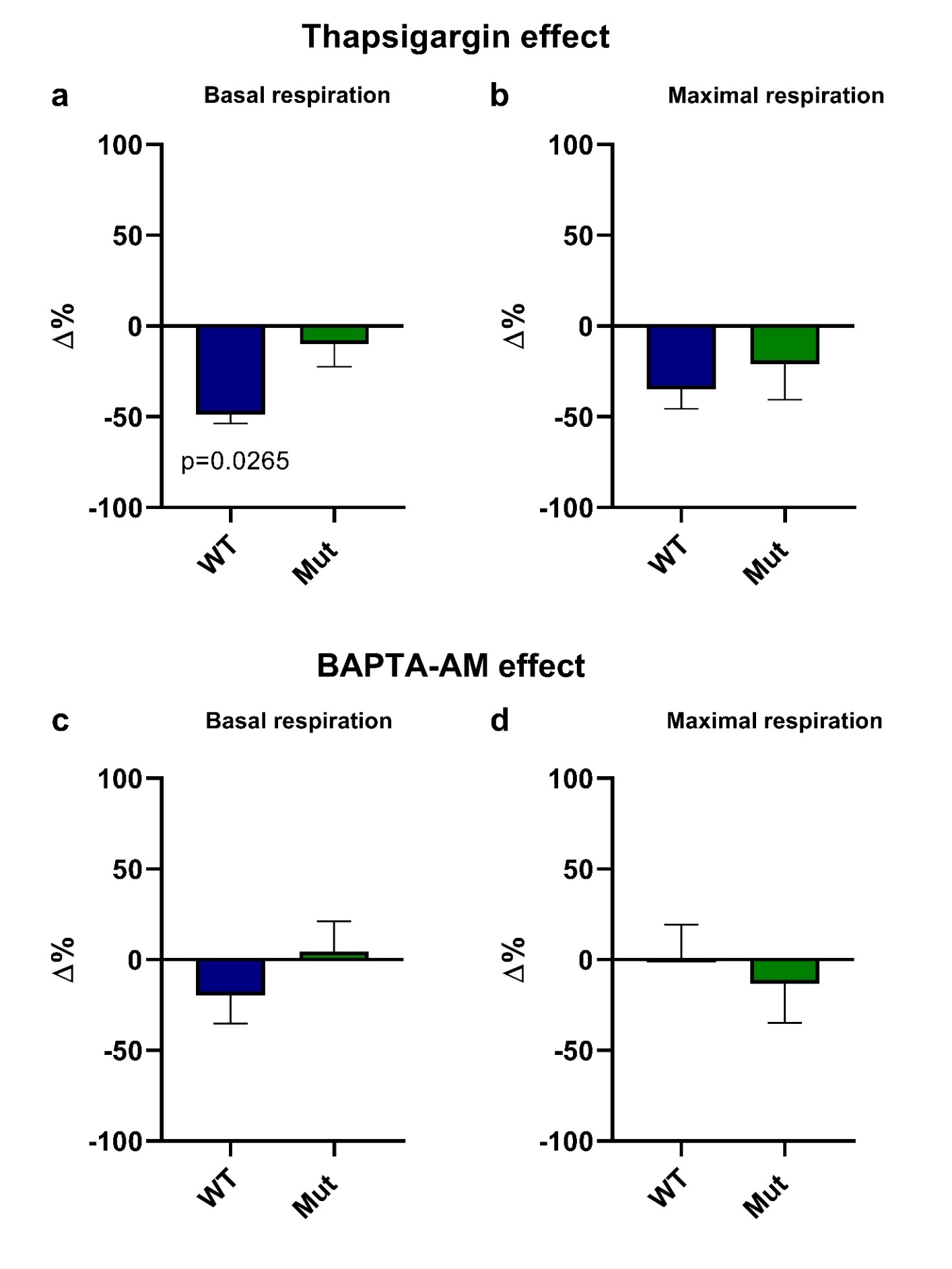

**Figure S6. Effect of BAPTA-AM and Thapsigargin on WT and Mut mitochondrial respiration** (**a**) Thapsigargin (THAPSI, 5 µM) effect on mitochondrial basal respiration (WT THAPSI: -49.51% ± 5.01; Mut THAPSI: -10.38% ± 12.65). (**b**) THAPSI effect on maximal respiration (WT THAPSI: -35.35% ± 10.68; Mut THAPSI: -21.30% ± 19.83). (**c**) BAPTA-AM (BAPTA, 5 µM) effect on mitochondrial basal respiration (WT BAPTA: -19.89% ± 15.79; Mut BAPTA: +4.60% ± 16.91). (**d**) BAPTA effect on mitochondrial maximal respiration (WT BAPTA: +0.54% ± 19.02; Mut BAPTA: -13.79% ± 21.75). WT THAPSI: n=18, N=4; WT BAPTA: n=12, N=3; Mut THAPSI: n=17, N=4; Mut BAPTA: n=11, N=3. Data are expressed as mean ± SEM. Kruskal-Wallis test.

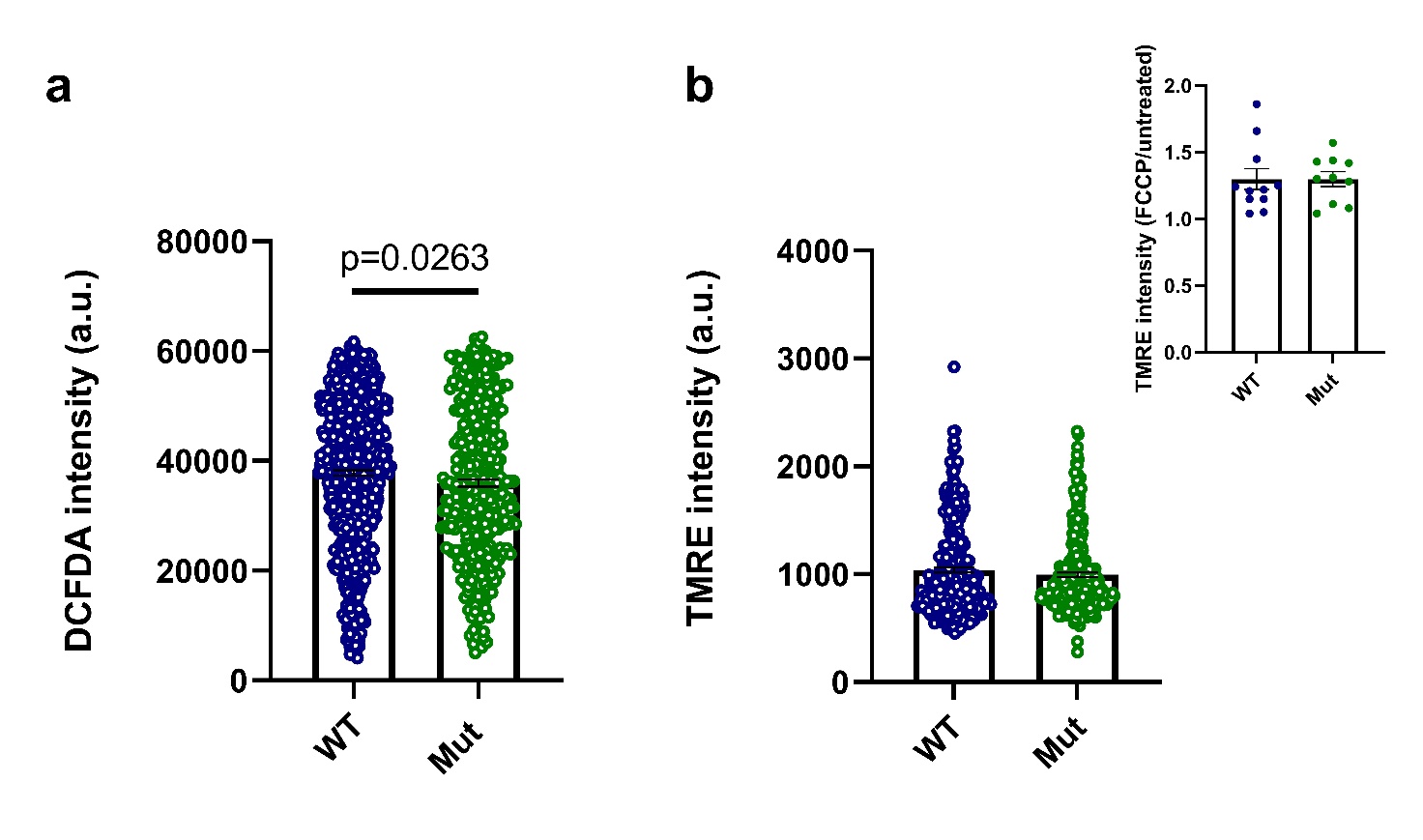

**Figure S7. ROS quantification and evaluation of** Ψ_m_ **in WT and Mut.** (**a**) DCFDA Fluorescence intensity (WT: 37827 ± 500, n=761, N=4; Mut: 35969 ± 672, n=430, N=4). (**b**) TMRE fluorescence intensity (WT: 1039 ± 24, n=311, N=4; Mut: 994 ± 21, n=305, N=4). **Inset**: TMRE fluorescence intensity after the incubation with FCCP normalized to fluorescence measured in control conditions (WT: 1.29 ± 0.07, n=11 wells, N=4; Mut: 1.29 ± 0.05, n=10 wells, N=4). Data are expressed as mean ± SEM. Unpaired Student’s t-test.

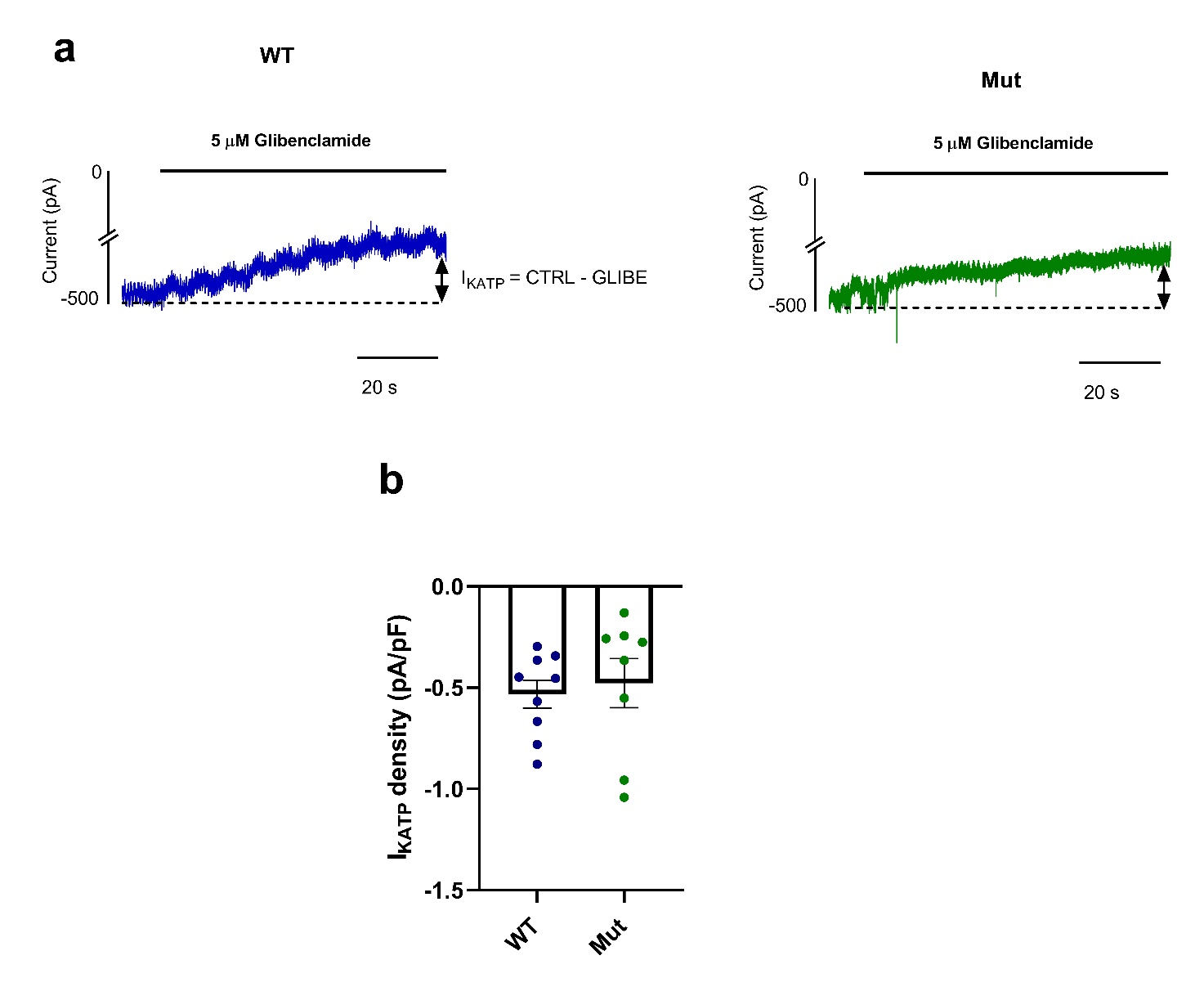

**Figure S8. Quantification of ATP-sensitive K^+^ current in WT and Mut.** (**a**) Representative current traces recorded with perforated patch technique at a holding potential of -120 mV, in control conditions and during perfusion with 5μM glibenclamide. (**b**) I_KATP_ density calculated as the current difference (control - glibe) normalized to membrane capacitance (WT: -0.53 ± 0.07, n=9; Mut: -0.47 ± 0.12, n=8). WT: N=7; Mut: N=6. Data are expressed as mean ± SEM. Unpaired Student’s t-test.

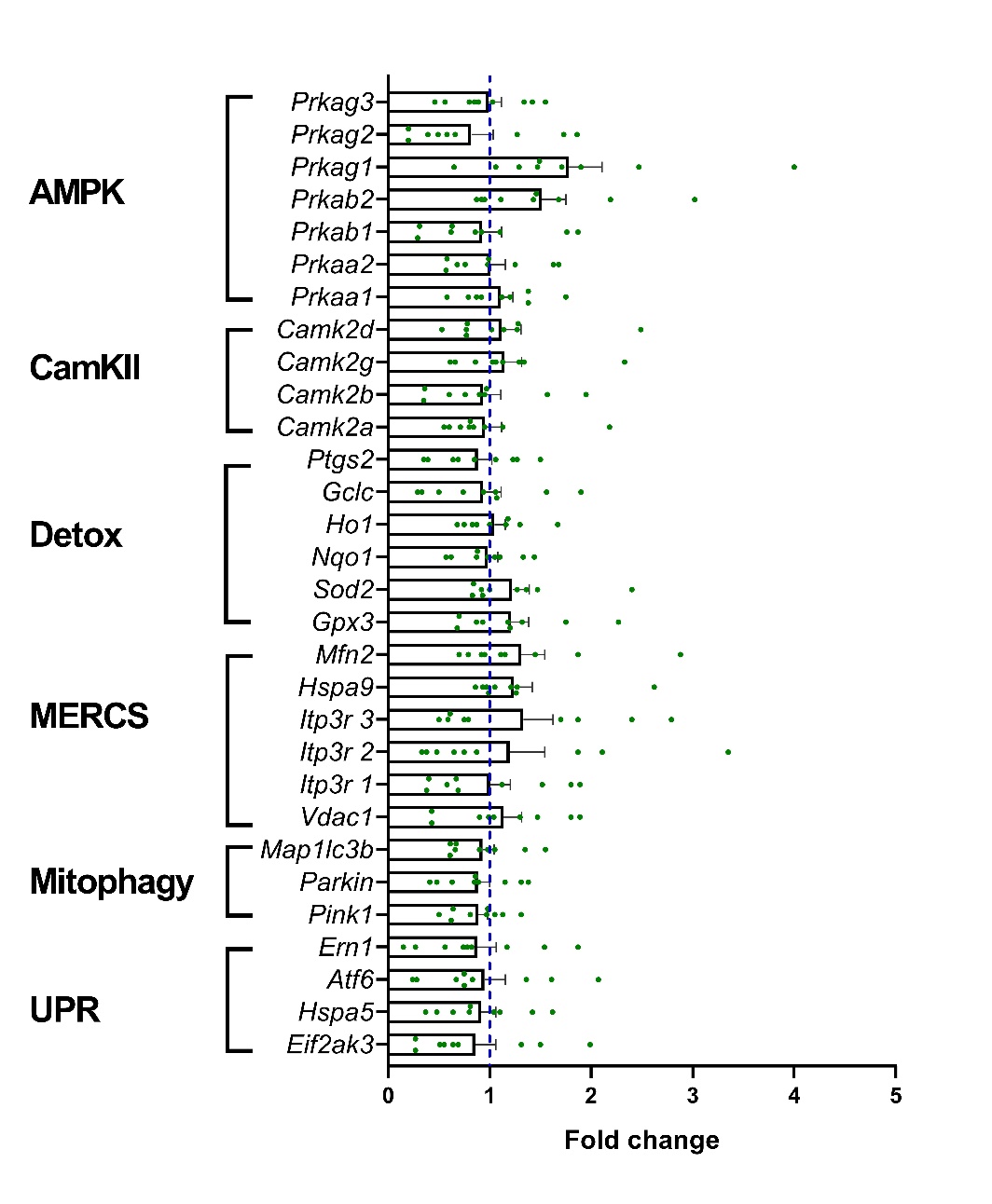

**Figure S9. Transcript analysis in WT and Mut.** Expression of AMPK (*Prkaa1*, *Prkaa2*, *Prkab1*, *Prkab2*, *Prkag1*, *Prkag2*, *Prkag3*), CaMKII isoforms (*Camk2a*, *Camk2b*, *Camk2g*, *Camk2d*) and genes associated to detoxification (*Ptgs2*, *Gclc*, *Ho1*, *Nqo1*, *Sod2*, *Gpx3*), mitochondria-ER contact sites (MERCS; *Mfn2*, *Hspa9*, *Itp3r1*, *Itp3r2*, *Itp3r3*, *Vdac1*), mitophagy (*Map1lc3b*, *Parkin*, *Pink1*) and unfolded protein response (UPR; *Ern1*, *Atf6*, *Hspa5*, *Eif2ak3*) in total RNA extracts of RV bioptic samples from WT and Mut mice. GAPDH was used as house-keeping gene and qRT-PCR data is shown as the fold change of target gene expression in Mut mice respect to WT mice (dotted blue line). Unpaired Student’s t-test.
